# Boosting the detection of enhancer-promoter loops via novel normalization methods for chromatin interaction data

**DOI:** 10.1101/2024.12.30.630839

**Authors:** Xiaotao Wang, Detong Shi, Feiyang Xue, Yunxia Liu, Hongbo Yang, Linghan Jiang

## Abstract

Accurately detecting enhancer-promoter loops from genome-wide interaction data, such as Hi-C, is crucial for understanding gene regulation. Current normalization methods, such as Iterative Correction and Eigenvector decomposition (ICE), are commonly used to remove biases in Hi-C data prior to chromatin loop detection. However, while structural or CTCF-associated loop signals are retained, enhancer-promoter interaction signals are often greatly diminished after ICE normalization and similar methods, making these regulatory loops harder to detect. To address this limitation, we developed Raichu, a novel method for normalizing chromatin contact data. Raichu identifies nearly twice as many chromatin loops as ICE, recovering almost all loops detected by ICE and revealing thousands of additional enhancer-promoter loops missed by ICE. With its enhanced sensitivity for regulatory loops, Raichu detects more biologically meaningful differential loops between conditions in the same cell type. Furthermore, Raichu performs consistently across different sequencing depths and platforms, including Hi-C, HiChIP, and single-cell Hi-C, making it a versatile tool for uncovering new insights into three-dimensional (3D) genomic organization and transcriptional regulation.

## Introduction

The spatial organization of chromatin plays a critical role in the precise control of transcriptional programs in mammalian cells^1, 2^. At kilobase to megabase scales, distant loci on the linear genome can come into close proximity in 3D space, forming structures known as chromatin loops^3^. These loops are broadly categorized into two types: structural loops, which connect CTCF-bound insulators, and regulatory loops, which link promoters to distal cis-regulatory elements, such as enhancers^4, 5^. The disruption or rewiring of these chromatin loops has been implicated in various developmental disorders and cancers^6, 7^.

A suite of experimental techniques has been developed to explore 3D genome organization and map chromatin loops^8^. Among these, Hi-C has become one of the most widely used methods due to its ability to capture chromatin contacts between all possible pairs of genomic loci^3, 9^. However, Hi-C data are subject to substantial biases, including sequence mappability, GC content, and restriction fragment length^10^. To mitigate these biases, a variety of computational normalization methods have been developed, which can be generally categorized into three groups: explicit methods, such as HiCNorm^11^ and OneD^12^, which model known sources of bias; implicit methods, such as ICE (Iterative Correction and Eigenvector Decomposition)^13^ and KR (Knight-Ruiz matrix balancing)^3^, which adjust the data without explicitly modeling biases; and hybrid methods, which combine features of both approaches^14^. Implicit methods like ICE and KR, owing to their simplicity and general applicability, have become the de facto standards for normalizing Hi-C data and are widely used in downstream analyses, including the identification of chromatin loops.

However, both ICE and KR have notable limitations. In a recent study by the 4D Nucleome Consortium, we observed that while these methods perform well in identifying CTCF-mediated loops, chromatin compartments, and other higher-order structures, they struggle to detect transcription-related loops^15^. This limitation arises from a key assumption in ICE and KR: that all genomic loci should have equal visibility in a Hi-C map. This assumption often leads to over-correction of interaction signals, causing low- frequency interactions, such as enhancer-promoter loops, to be under-detected despite their central role in gene regulation.

To address this long-standing gap in 3D genome analysis, we introduce Raichu, a novel computational method for normalizing chromatin contact data. Raichu is an implicit method that shares the simplicity of ICE and KR but diverges from their assumption of uniform visibility across genomic loci. Instead, Raichu employs an optimization-based approach that adjusts for variable interaction biases embedded in the raw data, enabling it to retain signals from biologically important, yet often subtle, chromatin interactions. Our results show that Raichu detects nearly twice as many chromatin loops as ICE, with a notable enrichment for enhancer-promoter loops that are critical for gene regulation. Importantly, Raichu outperforms existing methods in identifying differential loops between experimental conditions, providing new insights into how chromatin architecture regulates transcription in different cellular states.

Furthermore, Raichu demonstrates robustness across varying sequencing depths and 3D genomic platforms, making it a versatile tool for analyzing chromatin interactions.

## Results

### Limitations of existing Hi-C data normalization methods in detecting transcription-related loops

In this study, we focus on implicit normalization methods for Hi-C data, as explicit and hybrid methods require additional external data (e.g., mappability, GC content, fragment length), which limits their applicability to new genome assemblies where such data may be unavailable. Specifically, we compare two widely used matrix balancing implementations: Iterative Correction and Eigenvector decomposition (ICE), as implemented in the cooler package^16^, and Knight-Ruiz balancing (KR), available in the juicer toolkit^17^. Both cooler and juicer are central to the 3D genomic data analysis ecosystem and have been integrated as default Hi-C data processing tools by the 4D Nucleome Consortium^18^. While other software based on implicit methods also exists, they either implement similar matrix balancing algorithms or are less suited for standard downstream analysis workflows.

In Supplementary Figure 1, we present raw Hi-C contact maps alongside ICE- and KR-normalized maps for selected regions in GM12878 cells. As expected, ICE and KR normalization reduce noise, resulting in clearer visualization of topologically associating domains (TADs)^19, 20^ and chromatin loop structures. However, a closer examination reveals that many enhancer-involved loops (marked by H3K27ac peaks) are prominent in the raw Hi-C data but become significantly diminished following ICE or KR normalization, often to a level indistinguishable from background noise. This observation highlights the limitations of current normalization methods in preserving transcription-related interactions.

### Raichu: A novel computational method for normalizing chromatin contact data

Here, we present Raichu, a novel computational method for normalizing chromatin contact data. Raichu operates under the assumption that raw interaction intensities can be decomposed into two components: one representing the average interaction frequency as a function of genomic distance (accounting for the distance-dependent decay of Hi-C signals), and another representing loci-specific biases (Fig. 1a). Based on this assumption, Raichu employs an efficient optimization algorithm, leveraging dual annealing, to estimate a genome-wide bias vector that best explains the raw contact signals (Methods). This bias vector is then used to correct Hi-C data in a manner similar to ICE and KR.

**Figure 1.**
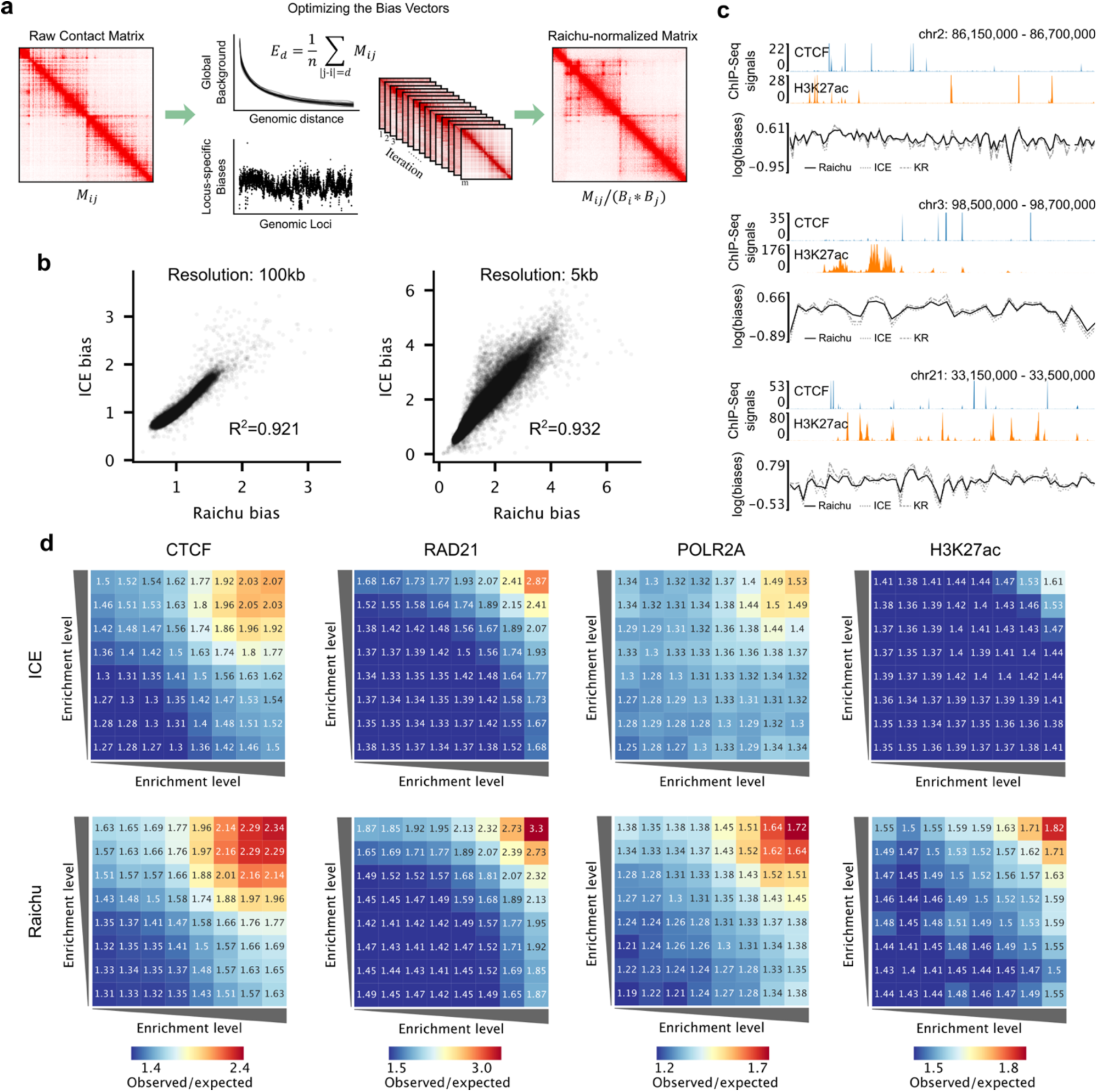
A novel computational method for normalizing chromatin contact data. **a**, Workflow of Raichu. **b**, Comparison of bias vectors calculated by ICE and Raichu at 100kb and 5kb resolutions. **c**, Example regions showing CTCF and H3K27ac ChIP-seq signals alongside bias vectors calculated by ICE, KR, and Raichu. While the bias vector calculated by Raichu is generally similar to those from ICE and KR, it exhibits lower variance. **d**, Comparison of average (observed/expected) contact signals between binding peaks of the indicated transcription factors or histone modifications for ICE and Raichu. Binding peaks were ranked by ChIP-seq signal enrichment and divided into 8 groups, and contact signals were averaged between pairs of groups.

To benchmark Raichu’s performance, we used the GM12878 Hi-C dataset^3^, which is one of the most deeply sequenced Hi-C datasets to date, and has extensive accompanying epigenomic and transcriptomic data in the same cell line. We first compared the bias vectors calculated by Raichu, ICE, and KR at various resolutions (Fig. 1b and Supplementary Fig. 2a). We observed a high concordance between the bias vectors produced by all three methods. Consequently, Raichu-normalized Hi-C signals were generally comparable to ICE/KR-normalized signals (Supplementary Fig. 2b). Furthermore, key chromatin features, such as compartments (measured by the first principal component, PC1^21^) and TADs (measured by insulation scores^21^), also demonstrated a high degree of concordance across Raichu, ICE, and KR (Supplementary Figs. 2c-d).

Upon closer inspection of specific genomic regions, however, we noticed that while Raichu’s bias vectors followed the same general trends as those of ICE and KR, the absolute values at the peaks and valleys were generally lower for Raichu (Fig. 1c). This difference translated into a stronger enrichment of Raichu-normalized signals for interactions that are weaker than canonical loop dots, yet clearly visible in the raw Hi-C data (highlighted by black circles in the bottom panel of Supplementary Fig. 2b). Given that various transcription factors (TFs) and histone modifications have been associated with chromatin loop formation, we next evaluated the enrichment of normalized contact signals between ChIP-seq peaks for selected TFs and histone modifications (Fig. 1d and Supplementary Fig. 3). Across all evaluated factors, Raichu-normalized signals consistently showed greater enrichment compared to ICE and KR, suggesting that Raichu enhances the detection of chromatin loops and may be better suited for capturing transcription-related interactions.

### Raichu identifies thousands of transcription-related loops missed by existing methods

To assess the effectiveness of Raichu in detecting chromatin loops, we applied HiCCUPS^3^, a widely used loop-calling algorithm, to ICE-normalized and Raichu-normalized contact maps at 5kb and 10kb resolutions (Methods). Strikingly, while ICE detected only 15,446 loops in GM12878 cells, Raichu identified 28,986 loops. Furthermore, 90.6% of loops detected by ICE (13,997 out of 15,446) were also identified by Raichu, whereas 51.7% of loops detected by Raichu (14,989 out of 28,986) were missed by ICE (Fig. 2a).

**Figure 2.**
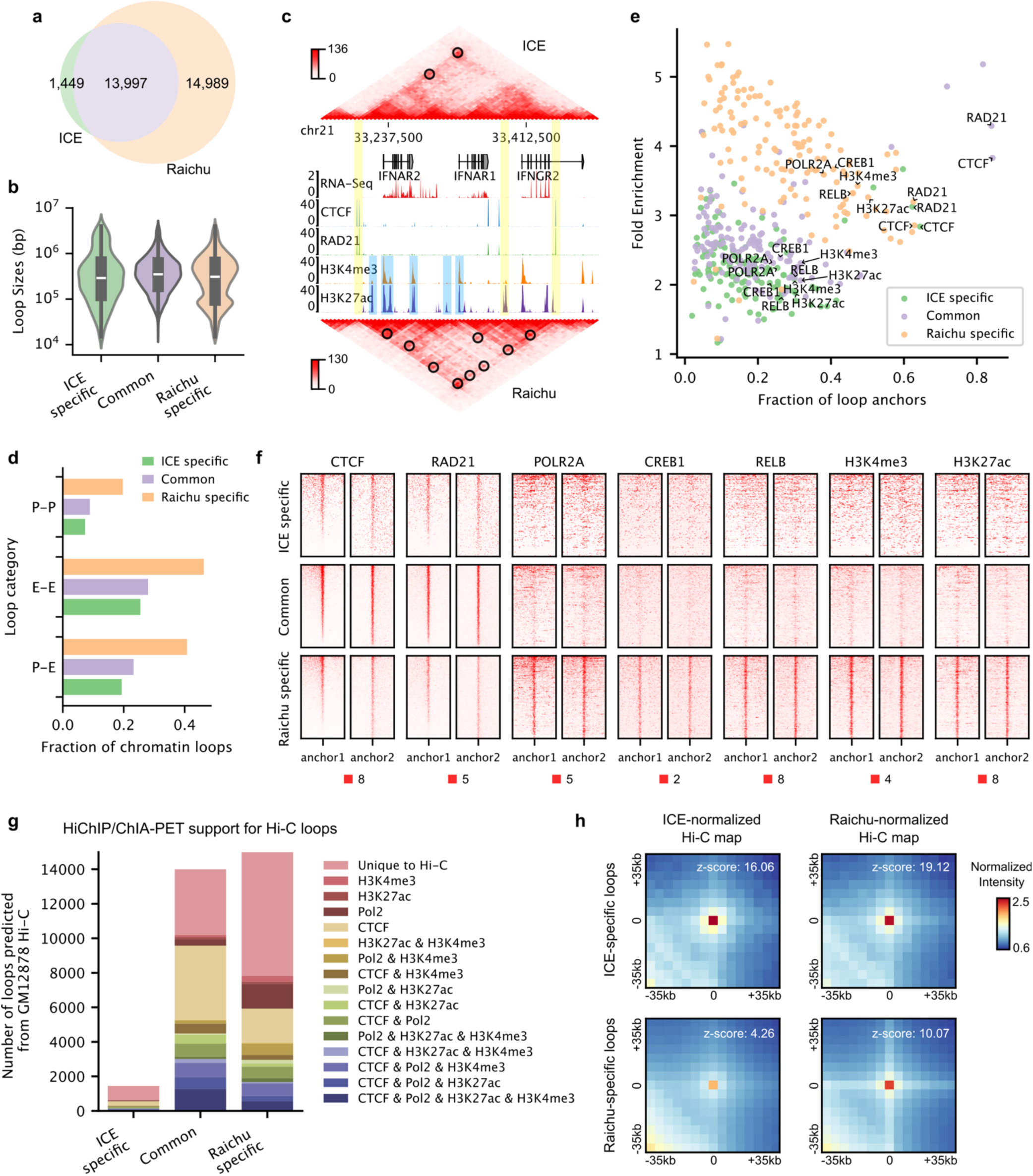
Raichu identifies thousands of transcription-related loops missed by ICE. HiCCUPS was used for loop detection. **a,** Venn diagram showing the overlap of loops detected by ICE and Raichu in GM12878 cells. **b**, Violin plots comparing the sizes of ICE-specific, Raichu-specific, and common loops detected by both ICE and Raichu. **c**, Comparison of contact signals for an example region between ICE and Raichu. Contact heatmaps are shown alongside gene annotations, RNA-seq data, and ChIP-seq signals for selected transcription factors and histone modifications. Black circles mark identified loops, yellow bars indicate loop anchors detected by both ICE and Raichu, and blue bars denote Raichu-specific loop anchors. **d**, Percentage of promoter-promoter (P-P), enhancer-enhancer (E-E), and promoter-enhancer (P-E) loops for ICE-specific, Raichu-specific, and common loops. **e,** Fraction of loop anchors bound versus fold enrichment for 132 transcription factors and 10 histone modifications, with different colors representing different loop categories. **f**, ChIP-seq binding profiles of selected transcription factors and histone modifications around both loop anchors for different loop categories, with each row representing one loop. **g**, Overlap with orthogonal ChIA-PET/HiChIP interactions for different loop categories. **h**, APA plots for ICE-specific and Raichu-specific loops in ICE- normalized and Raichu-normalized Hi-C maps.

We then classified the detected loops into three categories: ICE-specific loops, Raichu-specific loops, and common loops detected by both methods. While the average genomic distance between loop anchors was similar across these categories, Raichu-specific loops exhibited a distinct bimodal size distribution, suggesting that a subset of these loops may be transcription-related, given that transcription-related loops are often shorter in range compared to CTCF-mediated loops^5, 15, 22, 23^ (Fig. 2b).

To investigate the functional implications of these loops, we analyzed their overlap with ChIP-seq peaks for selected TFs and histone modifications. Figure 2c highlights an example region where ICE detected only two loops, both associated with CTCF/RAD21 ChIP-seq peaks. In comparison, Raichu not only identified both ICE-detected loops but also revealed seven additional loops in the same region, most of which overlapped with either H3K27ac peaks (marking active enhancers and promoters) or H3K4me3 peaks (marking promoters) at both anchors, suggesting their involvement in transcriptional regulation. Genome-wide, Raichu-specific loops showed a higher proportion of enhancer-promoter (E-P), enhancer- enhancer (E-E), and promoter-promoter (P-P) interactions (40.8%, 46.3%, and 19.7%, respectively) compared to ICE-specific loops (19.3%, 25.5%, and 7.3%) and common loops (23.2%, 28.0%, and 8.9%). To systematically examine the factors associated with these loops, we calculated the fold enrichment of 132 TFs and 10 histone modifications at loop anchors using ENCODE data. As expected, common loops showed the highest enrichment for CTCF and RAD21. While ICE-specific and Raichu-specific loops exhibited comparable enrichment for CTCF and RAD21, Raichu-specific loops demonstrated substantially greater enrichment for a broader range of TFs and histone modifications closely associated with transcriptional regulation, including RNA polymerase II (POLR2A), CREB1, RELB, H3K4me3, and H3K27ac (Figs. 2e-f and Supplementary Fig. 4).

To further explore the nature of loops detected by ICE and Raichu, we validated them against four orthogonal 3D genomic datasets (Supplementary Table 2): CTCF ChIA-PET (targeting CTCF-mediated interactions), Pol2 ChIA-PET (targeting RNA polymerase II-mediated interactions), H3K27ac HiChIP (targeting H3K27ac interactions), and H3K4me3 HiChIP (targeting H3K4me3 interactions). As expected, common loops exhibited the highest validation rate across these datasets (72.8%; 10,191 out of 13,997 common loops), followed by Raichu-specific loops (52.2%; 7,823 out of 14,989) and ICE-specific loops (44.2%; 640 out of 1,449) (Fig. 2g). Notably, among the validated Raichu-specific loops, 74.6% (5,837 out of 7,823) overlapped with transcription-related interactions identified by Pol2 ChIA-PET, H3K27ac HiChIP, or H3K4me3 HiChIP, compared to 60.5% for ICE-specific loops and 57.8% for common loops.

It is worth noting that a loop’s absence from a call set does not necessarily indicate a lack of enrichment; it may simply fall below the statistical threshold. To evaluate whether ICE-specific and Raichu-specific loops were enriched in different normalized Hi-C maps, we performed an Aggregate Peak Analysis (APA) (Fig. 2h). Interestingly, ICE-specific loops showed even greater enrichment in Raichu-normalized maps than in ICE-normalized maps, indicating that these loops are detectable by Raichu. Their absence likely stems from ranking below the threshold. In contrast, Raichu-specific loops displayed much weaker enrichment in ICE-normalized data, suggesting they are genuinely missed by ICE, even with relaxed cutoffs.

We also benchmarked Raichu against KR normalization and evaluated its performance across different loop-calling algorithms^24^, with similar results observed in each comparison (Supplementary Figs. 5-6). Collectively, these findings demonstrate that Raichu surpasses existing methods in its ability to detect transcription-related loops.

### Raichu is robust across different sequencing depths

To evaulate the impact of sequencing depth on the performance of Raichu, we computationally down- sampled the GM12878 dataset to 23 different depths, ranging from 100 million to 2 billion intra-chromosomal reads (Methods). As expected, the number of detected loops decreased with reduced sequencing depth for both ICE and Raichu. Notably, Raichu consistently detected approximately twice as many loops as ICE across all tested depths (Figs. 3a-b). Consistent with results from the full GM12878 dataset, the additional loops detected by Raichu were specifically enriched for transcription-related interactions, as confirmed by orthogonal datasets (Fig. 3c).

**Figure 3.**
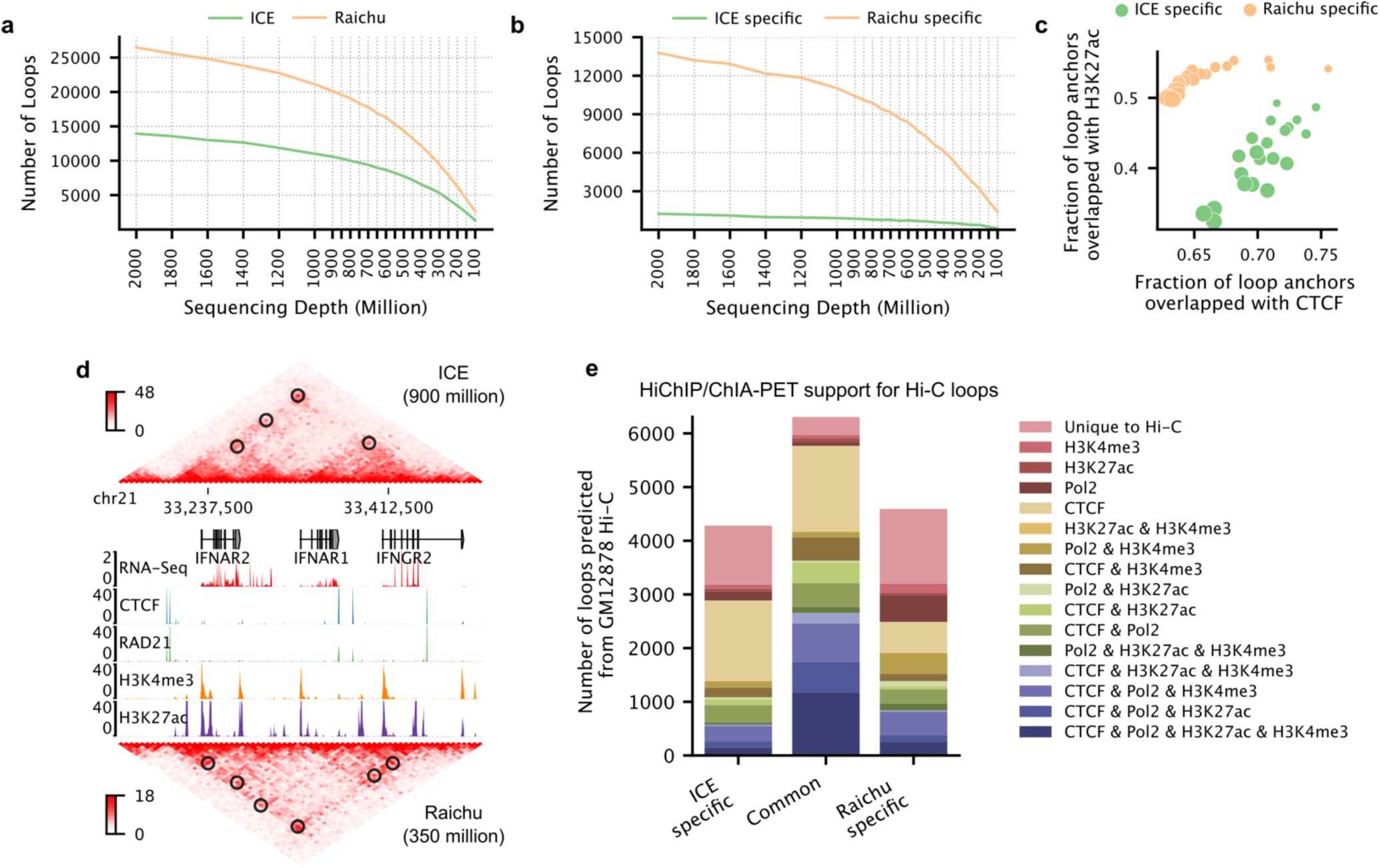
Raichu performs well across various sequencing depths. **a,** Number of loops detected by ICE and Raichu across a series of down-sampled GM12878 Hi-C maps. **b**, Counts of ICE-specific and Raichu- specific loops for the same series of down-sampled GM12878 Hi-C maps. **c**, Fraction of loop anchors overlapping with H3K27ac versus fraction overlapping with CTCF for ICE-specific and Raichu-specific loops across different down-sampling levels. Dot sizes represent sequencing depths. **d**, Comparison of loops identified by ICE with 900 million usable reads and Raichu with 350 million usable reads. The panel includes contact heatmaps, gene annotations, RNA-Seq data, and ChIP-Seq data for selected transcription factors and histone modifications. Black circles mark identified loops. **e**, Overlap with orthogonal ChIA-PET/HiChIP interactions for ICE-specific, Raichu-specific, and common loops, comparing loops detected by ICE with 900 million usable reads and Raichu with 350 million usable reads.

Raichu’s greater power in detecting chromatin loops enables it to achieve loop counts at lower sequencing depths that are comparable to those of ICE at higher sequencing depths (Fig. 3a). For example, while ICE detected 10,589 loops at a depth of ∼900 million usable reads, Raichu identified a similar 10,900 loops with only ∼350 million reads. Of these, 59.6% (6,307 out of 10,589) of loops detected by ICE overlapped with 57.9% (6,307 out of 10,900) of loops detected by Raichu. Importantly, validation rates by orthogonal ChIA-PET/HiChIP datasets were comparable between ICE-specific and Raichu- specific loops (74.3% vs. 69.6%) in this comparison. However, 82.2% of validated Raichu-specific loops overlapped with transcription-related interactions by Pol2 ChIA-PET, H3K27ac HiChIP, or H3K4me3 HiChIP datasets, compared to only 53.0% for ICE-specific loops (Figs. 3d-e). Similar trends were observed at additional sequencing depths (Supplementary Fig. 7).

Together, these findings demonstrate that Raichu consistently detects thousands of additional chromatin loops associated with gene regulation across a broad range of sequencing depths.

### Raichu enhances the detection of chromatin loops in single-cell Hi-C data

We next applied Raichu to a GM12878 single-cell Hi-C dataset containing 221 single cells, with a median depth of approximately 1.08 million contacts^25^. To compare the performance of Raichu and ICE on this dataset, we ranked all 221 cells by sequencing depth and created a series of pseudo-bulk datasets by pooling the top 130 deepest single cells (Supplementary Fig. 8a). Then for each pooled dataset, loops were identified using HiCCUPS with either ICE-normalized or Raichu-normalized contact maps as input, and the detected loops were merged across 5kb, 10kb, and 25kb resolutions (Methods).

As shown in Figure 4a, Raichu detected 1.24 to 1.75 times more loops than ICE. Even with as few as 5 cells (∼10 million contacts), Raichu identified 1,697 loops, whereas ICE detected only 970 loops on the same dataset. Using a list of interactions compiled from orthogonal 3D genomic datasets (including Hi-C, CTCF ChIA-PET, RNAPII ChIA-PET, CTCF HiChIP, H3K27ac HiChIP, SMC1A HiChIP, H3K4me3 PLAC-Seq, TrAC-Loop, and HiCAR) as true positive loops, we evaluated the precision and recall of Raichu and ICE across all pooled datasets derived from different cell counts (Supplementary Figs. 8b-c). Notably, Raichu achieved a higher F1 score compared to ICE across all tested cases (Fig. 4b).

**Figure 4.**
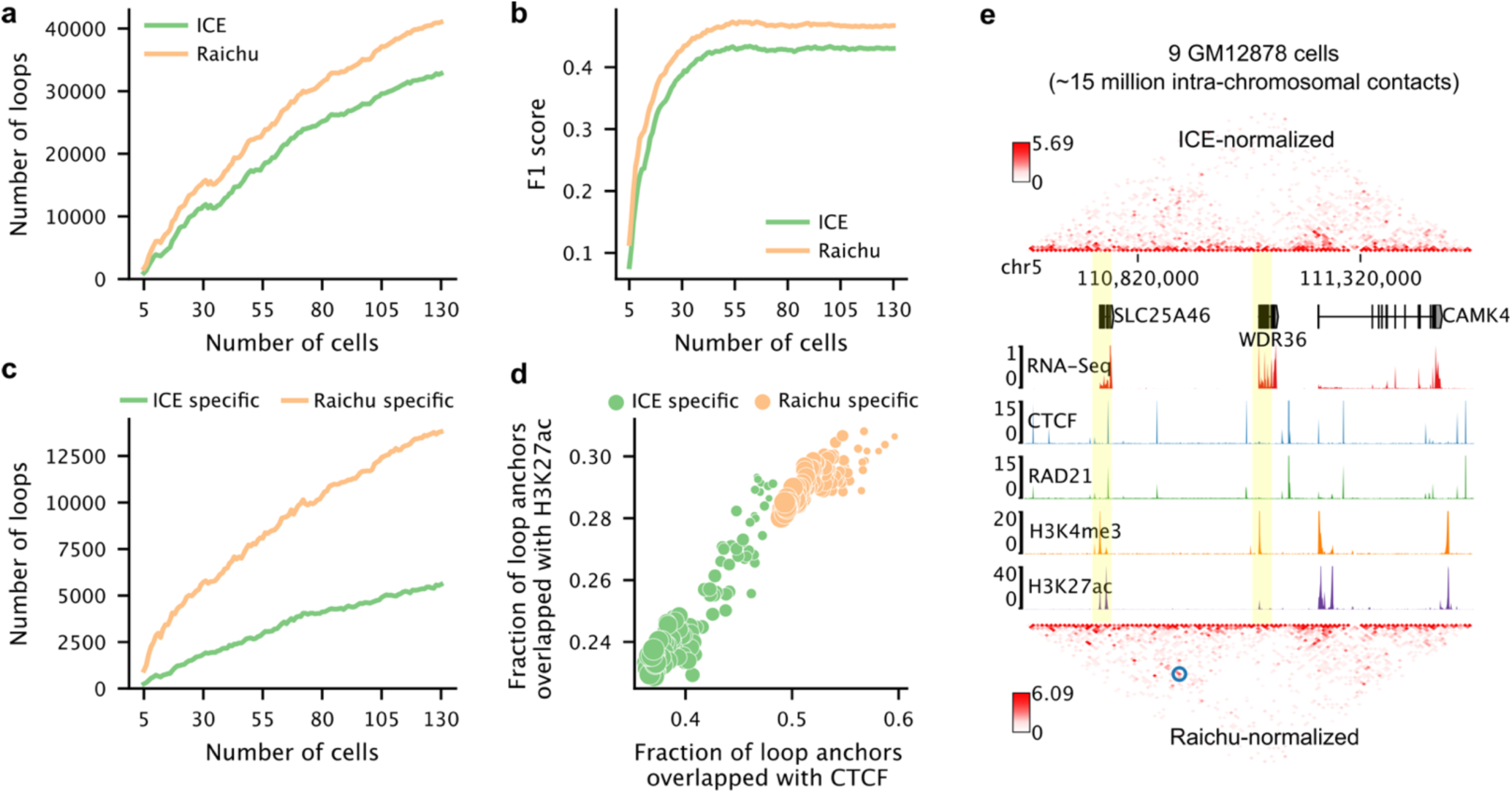
Raichu enhances the detection of chromatin loops in single-cell Hi-C data. **a,** Number of loops detected by ICE and Raichu as a function of the number of merged GM12878 cells. **b**, Comparison of F1 scores for loops detected by ICE and Raichu across the same merged datasets. **c**, Counts of ICE-specific and Raichu-specific loops identified across increasing numbers of merged GM12878 cells. **d**, Fraction of loop anchors overlapping with H3K27ac versus fraction overlapping with CTCF for ICE-specific and Raichu-specific loops. Dot sizes represent the number of cells. **e**, Comparison of contact signals and loop calls between ICE and Raichu for maps merged from 9 GM12878 cells. Blue circles mark identified loops, and yellow bars highlight the two loop anchors uniquely detected by Raichu-normalized signals.

For each pooled map, the number of Raichu-specific loops was 2.45 to 4.59 times greater than ICE- specific loops (Fig. 4c). More importantly, a higher percentage of loop anchors detected by Raichu overlapped with CTCF and H3K27ac binding compared to ICE-specific loops, indicating that Raichu excels in detecting both CTCF-related and transcription-related loops from single-cell Hi-C data (Figs. 4d-e).

### Raichu detects unique differential loops involved in transcriptional regulation

During cell development or drug treatment, chromatin looping structures often undergo dramatic changes, leading to gene activation or repression^26, 27, 28, 29^. Accurately detecting these changes between conditions is critical for understanding the molecular mechanisms of gene regulation underlying specific cell states.

To demonstrate the power of Raichu in detecting such changes, we utilized a cell system we constructed in a previous study^30^. Briefly, through targeted sequencing of 5,008 patients, we identified a key regulatory germline variant in the GATA3 enhancer that is closely associated with Philadelphia chromosome-like acute lymphoblastic leukemia. To study its role in 3D genome organization and gene regulation, we genetically engineered the wild-type C/C allele into the risk A/A allele in GM12878 cells. In wild-type cells, GATA3 is only moderately expressed. In engineered cells, however, the activity of the GATA3 enhancer is significantly increased, resulting in high GATA3 expression. Additionally, GATA3 binding is increased at thousands of enhancers, further driving the transcription of downstream genes.

In our original study, using ICE-normalized Hi-C maps, we identified only a few cases where differential chromatin loops were associated with gene regulation. Here, we explored whether the enhanced sensitivity of Raichu in detecting transcription-related loops could improve the detection of meaningful chromatin looping changes between wild-type (C/C) and engineered (A/A) cells.

Raichu indeed demonstrated clear improvements when we examined specific loci. For example, at the SUPT16H gene locus, ICE-normalized signals failed to reveal meaningful loop differences between wild- type and engineered cells (Fig. 5a, left). In contrast, Raichu identified an additional loop unique to the engineered cells, linking the SUPT16H gene — encoding SPT16, a subunit of the FACT (Facilitates Chromatin Transcription) complex critical for chromatin remodeling, transcriptional regulation, and genomic stability — to a downstream enhancer with specific GATA3 binding in engineered cells (Fig. 5a, right). This example, along with others (Supplementary Fig. 9), highlights the ability of Raichu in uncovering novel regulatory mechanisms during cell development and disease.

**Figure 5.**
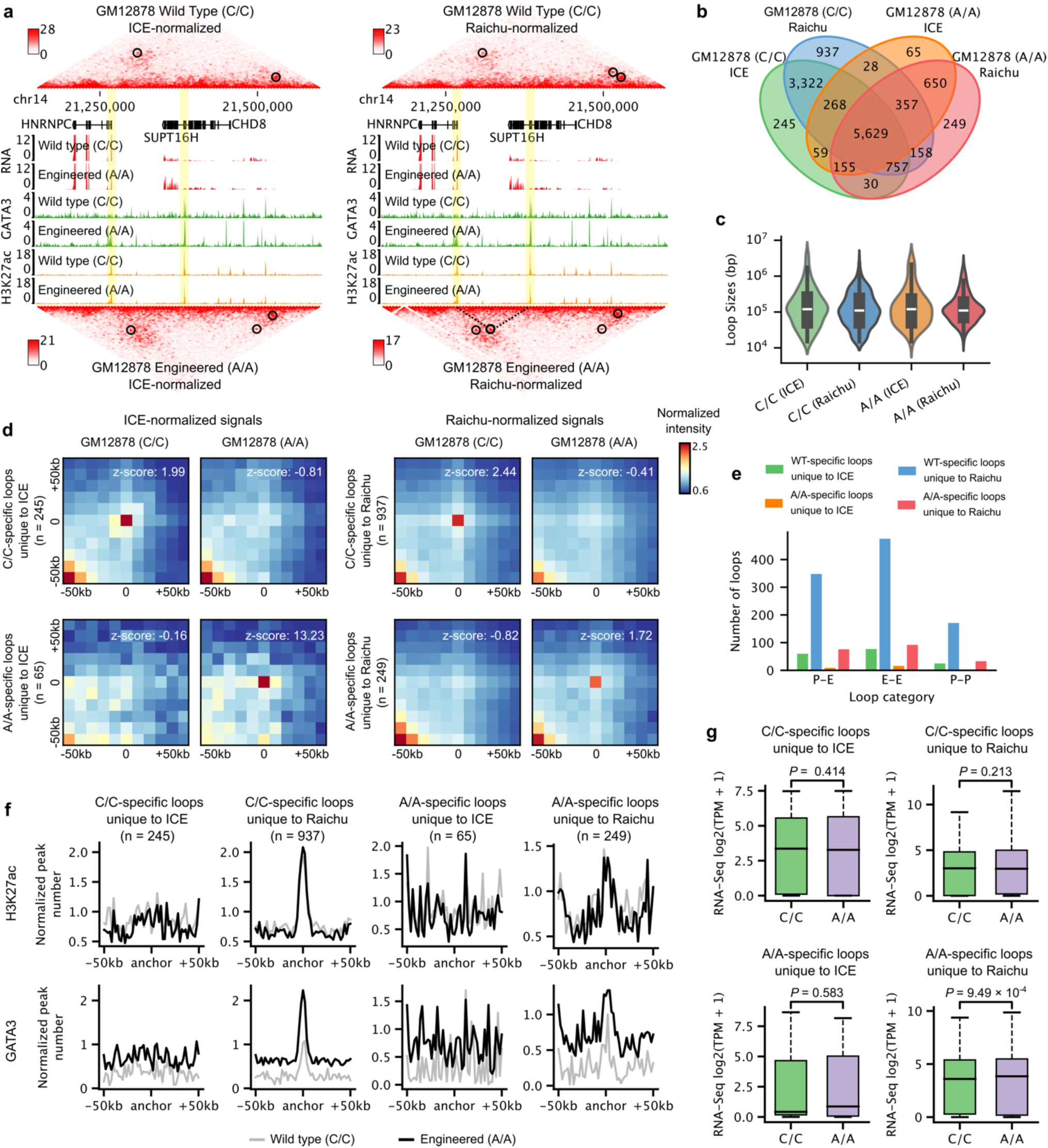
Raichu detects unique differential loops associated with transcriptional regulation. **a,** Comparison of Hi-C contact maps, RNA-Seq, GATA3 ChIP-Seq, and H3K27ac ChIP-Seq signals between wild-type (C/C) and engineered (A/A) GM12878 cells. The left panel displays ICE-normalized Hi-C signals, while the right panel shows Raichu-normalized signals. Black circles mark the identified loops, and the anchors of an A/A-specific loop unique to Raichu are highlighted in yellow. **b**, Venn diagram showing overlap between loops detected by ICE and Raichu in wild-type and engineered GM12878 cells. **c**, Loop size distributions for C/C-specific loops unique to ICE and Raichu, and A/A-specific loops unique to ICE and Raichu. **d**, APA plots for C/C-specific loops unique to ICE and Raichu, and A/A-specific loops unique to ICE and Raichu in C/C and A/A Hi-C maps. **e**, Counts of promoter-enhancer (P-E), enhancer-enhancer (E-E), and promoter-promoter (P-P) loops within the different loop categories as indicated. **f**, H3K27ac and GATA3 ChIP-Seq profiles surrounding loop anchors for the different loop categories as indicated. **g**, Comparison of quantile-normalized transcription levels between wild-type and engineered cells for target genes of GATA3-bound enhancers linked by the indicated loop categories.

Using Hi-C data normalized by either ICE or Raichu, we identified 4,504 wild-type (C/C)-specific loops. Of these, 937 were unique to Raichu, while only 245 were unique to ICE. Similarly, we identified 964 loops specific to the engineered cells, with 249 unique to Raichu and only 65 unique to ICE (Figs. 5b-d). APA plots confirmed that C/C-specific loops, whether unique to ICE or Raichu, were enriched in wild-type cells but depleted in engineered cells, and vice versa for A/A-specific loops (Fig. 5d; Methods). While the size distribution of differential loops was comparable across groups, Raichu-specific loops had shorter average genomic distances between anchors compared to ICE-specific loops (203kb vs. 298kb for C/C- specific loops; 202kb vs. 306kb for A/A-specific loops), consistent with Raichu’s higher sensitivity for short-range loops (Fig. 5c and Fig. 2b).

To associate these differential loops with genomic functions, we categorized them into P-E, E-E, and P-P loops. Strinkingly, Raichu-specific loops contained 5.75 – 16.5 times more of these transcription- related loops compared to ICE-specific loops (Fig. 5e). Furthermore, Raichu-specific differential loops showed significant enrichment for H3K27ac and GATA3 binding signals at their anchors, a pattern not observed for ICE-specific loops (Fig. 5f). Comparing transcription levels between wild-type and engineered cells revealed that only Raichu-unique A/A-specific loops were associated with significant gene upregulation in engineered cells (Fig. 5g).

Collectively, these results demonstrate that Raichu improves the detection of meaningful differential loops between conditions, capturing loops closely associated with gene regulation and cell function that ICE might miss.

### Raichu performs well on HiChIP data

In addition to Hi-C, numerous methods have been developed to study 3D genome organization. Some, such as Micro-C^31^ and DNA SPRITE^32^, aim to profile chromatin interactions across the genome without bias and are typically normalized using algorithms designed for Hi-C. Other methods, including HiChIP^33^, ^34^, ChIA-PET^35^, and Capture Hi-C^36^, are designed to enrich interactions mediated by specific proteins or DNA elements. These pulldown-based methods, however, often lack dedicated normalization approaches. Consequently, their data are either left unnormalized or processed using Hi-C-derived normalization methods, which may fail to address their unique biases.

To evaluate whether Raichu is applicable to these methods, we analyzed an H3K27ac HiChIP dataset from K562 cells^37^, which enriches interactions involving H3K27ac, a marker for active enhancers. First, we visually compared the raw, ICE-normalized, and Raichu-normalized contact maps. In the raw map, H3K27ac-enriched regions formed extensive chromatin stripes, with interactions along the stripes often spanning multiple TADs. In the ICE-normalized map, while most interactions were relatively confined within TADs, the H3K27ac-specific signals were almost entirely lost. In contrast, the Raichu-normalized map retained both TAD structures and H3K27ac-specific signals (Fig. 6a). Aggregation analysis of TADs further supported these observations (Fig. 6b). Since TADs are structural units where enhancers preferentially regulate genes within the same TAD and rarely interact with genes outside the TAD^19^, these results suggest that Raichu-normalized signals achieve a better balance of sensitivity and specificity for detecting enhancer-promoter interactions compared to raw or ICE-normalized signals.

**Figure 6.**
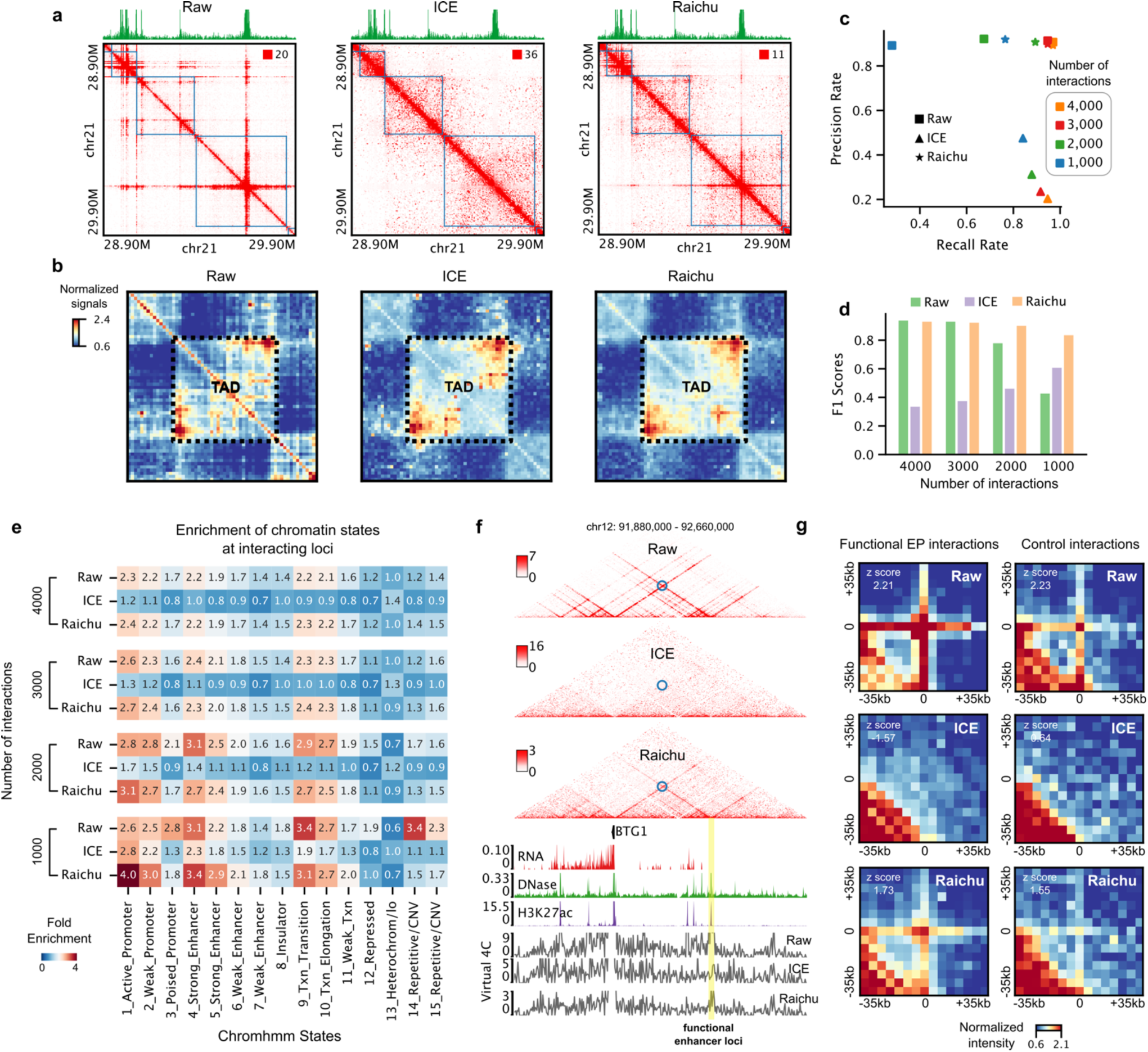
Application of Raichu in normalizing H3K27ac HiChIP data. **a,** Comparison of raw, ICE-normalized, and Raichu-normalized HiChIP signals for an example region in K562 cells. The top green track shows one-dimensional HiChIP coverage, and blue squares indicate TAD regions identified from Hi-C data in the same cell line. **b**, Aggregate plots of regions surrounding TADs using raw, ICE-normalized, and Raichu- normalized signals. **c**, Recall and precision rates of different normalization methods in detecting enhancer-associated loops. **d**, F1 scores of different normalization methods in detecting enhancer-associated loops. **e**, Fold enrichment of ChromHMM states at interacting loci identified using different normalization methods. **f**, HiChIP maps and supporting data for the BTG1 region, including RNA-Seq, DNase-Seq, H3K27ac ChIP-Seq, and virtual 4C tracks using the BTG1 promoter as the viewpoint. The yellow bar marks a functional enhancer validated by CRISPR as regulating BTG1 expression in K562 cells, and blue circles highlight the functional interaction between BTG1 and the enhancer locus. **g**, APA plots for functional and control enhancer-promoter interactions in K562 cells. Control interactions use the same enhancer loci as the functional interactions but pair them with loci at similar genomic distances on the opposite side.

We then performed loop detection on raw, ICE-normalized, and Raichu-normalized maps, selecting loops at various significance thresholds to mitigate cutoff effects (Methods). Given that K562 is a cancer cell line with numerous structural vairations (SVs), we focused on chromosome 21, which is relatively free of SVs in this cell line. Consistent with previous observations, most loops detected using raw and Raichu-normalized signals were associated with H3K27ac-enriched regions. In constrast, loops detected using ICE-normalized signals frequently linked non-specific loci, likely reflecting artifacts introduced during normalization. Notably, at stricter thresholds (top 2,000 and 1,000 interactions), certain regions with visibly enriched signals in the raw maps showed no loops. However, these regions consistently displayed loops in Raichu-normalized maps across all thresholds (Supplementary Fig. 10).

To systematically evaluate the specificity and sensitivity of raw, ICE-normalized, and Raichu- normalized maps for detecting enhancer-associated loops, we defined a high-confidence set of H3K27ac- enriched regions by integrating H3K27ac HiChIP and ChIP-Seq data (Methods). Approximately 90% of loops detected using raw and Raichu-normalized signals overlapped with H3K27ac-enriched regions at either anchor (precision), compared to less than 50% for ICE-normalized loops (Fig. 6c). Recall rates, representing the percentage of H3K27ac-enriched regions overlapping with loop anchors, were similar for raw and Raichu-normalized signals at more relaxed thresholds (top 4,000 and 3,000 interactions). However, recall rates for Raichu-normalized loops declined more slowly at stricter thresholds, resulting in higher F1 scores, particularly under stringent p-value cutoffs (Fig. 6d). Raichu-normalized loops also showed greater enrichment for active promoters and enhancers compared to raw and ICE-normalized loops. Interestingly, loops from raw maps were disproportionately enriched for repetitive or copy number variation (CNV) regions, highlighting the importance of correcting biases inherent in raw HiChIP data (Fig. 6e).

Since H3K27ac HiChIP is primarily used to identify loops involved in gene regulation, we next examined how functional interactions validated by CRISPR perturbation experiments^38^ were enriched in maps normalized by different methods. Nearly all functional interactions were depleted in ICE-normalized maps, whereas many were enriched in raw and Raichu-normalized maps (Fig. 6f and Supplementary Fig. 11). Interestingly, Raichu-normalized maps exhibited fewer stripe signals compared to raw maps, resulting in slightly higher enrichment of functional interactions relative to control interactions (z-scores 1.73 vs. 1.55), which were indistinguishable in raw maps (z-scores 2.21 vs. 2.23) (Fig. 6g).

Collectively, these results demonstrate that Raichu outperforms both unnormalized and ICE- normalized signals in detecting enhancer-associated loops and functional enhancer-promoter interactions from H3K27ac HiChIP data.

## Discussion

Recent years have highlighted the critical role of chromatin organization in transcriptional regulation, yet existing normalization methods, such as ICE, have struggled to capture transcription-related chromatin loops from Hi-C data. Here, we present Raichu, a novel computational method for normalizing chromatin contact data. Unlike matrix balancing methods, Raichu does not assume uniform visibility across all genomic loci. Instead, it decomposes the expected frequency of each contact into two components: 1) a global background that accounts for the distance-dependent decay of interaction signals, and 2) locus- specific biases. An effcient optimization algorithm is then used to calculate the bias vector. We demonstrate that Raichu avoids the over-correction issues inherent to ICE, notably enhancing the ability of Hi-C to detect transcription-related loops. Furthermore, Raichu performs robustly across varying sequencing depths and is compatible with other 3D genomic platforms, underscoring its versatility.

In this study, we compared Raichu with two of the most popular implementations of the matrix balancing algorithm, ICE and KR. While other methods, such as HiCNorm^11^, OneD^12^, and HiCorr^14^, normalize Hi-C data by explicitly modeling known genomic biases, we excluded them from the comparison because they require external reference data, which are typically pre-built for only a limited number of reference genomes. Built on the cooler Python package^16^, Raichu does not require any external data and stores the calculated bias vector in the same format as ICE, ensuring seamless compatibility with downstream tools for analyzing compartments, TADs, loops, and other features. Additionally, Raichu is computationally efficient: running on a Hi-C dataset with 4 billion usable reads at 5kb resolution with 4 CPU cores, it completed the calculation in approximately 4 hours, requiring only 17 GB of memory (Supplementary Table 1). These features make Raichu a powerful tool that can serve as a replacement for ICE.

Recent advances in single-cell Hi-C and related techniques have significantly enhanced our ability to study chromatin organization at the resolution of individual cells. These innovations now enable the simultaneous profiling of chromatin contacts alongside additional layers of epigenomic and transcriptomic information, such as DNA methylation^29, 39, 40^, RNA expression^25, 41, 42^, and chromatin accessibility (ATAC)^43^. Such integrated approaches make it possible to reliably identify cell groups or cell types and investigate cell-type-specific chromatin contacts and their roles in gene regulation. Typically, the number of cells in each identified group ranges from a few dozen to a few hundred. Using a recently published single-cell Hi-C dataset in GM12878 cells, we demonstrated that Raichu is applicable to datasets with as few as 5 cells, representing approximately 10 million chromatin contacts. Even under these sparse conditions, Raichu outperformed existing Hi-C normalization methods in detecting both CTCF-related and transcription-related loops (Fig. 4). This result highlights Raichu’s robustness and utility in handling sparse datasets characteristic of single-cell Hi-C experiments. Future applications of Raichu to pseudo- bulk Hi-C contact maps derived from single-cell multi-omics techniques could provide researchers with new insights into the relationship between gene regulation and cell-type-specific 3D genome organization.

The close association between 3D genome organization and transcription regulation has long been recognized, with disruptions in chromatin architecture linked to developmental diseases and cancers. However, the relationship between chromatin loops and gene regulation remains elusive, particularly in conditions with minimal changes in cell state, such as drug treatments or heat shock. In these scenarios, studies using ICE often report negligible chromatin loop changes, reinforcing the belief that most loops are pre-established^44, 45^. Our analysis reveals that transcription-related loops are frequently attenuated in ICE-normalized data, suggesting that prior studies may have underestimated 3D genomic changes. By applying Raichu to an engineered GM12878 cell system, we identified significant differential loops closely associated with gene transcription that ICE failed to detect. These findings highlight Raichu’s ability to uncover novel insights into the interplay between 3D genome organization and gene regulation.

Future applications of Raichu may redefine how we study 3D genomic changes, offering a more accurate perspective on their roles in transcriptional regulation and cellular function. This capability is particularly promising for elucidating subtle chromatin dynamics in development, disease progression, and therapeutic interventions. By addressing the limitations of existing normalization methods, Raichu opens new avenues for the exploring chromatin architecture and its impact on gene expression.

## Methods

### Workflow of Raichu

The core rationale underlying Raichu is that each contact in the matrix can be decomposed into two components: (1) global background (*E_d_*), which accounts for the distance-dependent decay of interaction signals (i.e., genomic loci in close proximity are more likely to interact than those farther apart), and (2) locus-specific biases (*B*), which capture variations in interaction frequency attributable to individual loci. Once these biases are calculated, contact counts can be normalized in a manner similar to ICE and other implicit normalization methods for chromatin contact data.

Given an intra-chromosomal contact matrix *M*, the global background is calculated as:

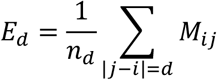

where *n_d_* represents the number of bin pairs separated by distance *d*, and *M_ij_* represents the observed interaction frequency between bin *i* and bin *j*.

Assuming locus-specific biases are multiplicative, each contact *M_ij_* is modeled as:

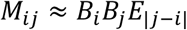

The objective is to estimate *B* such that the product *B_i_B_j_E*_|*j–i*|_ approximates the observed contact count *M_ij_*. This is formulated as the following optimization problem:

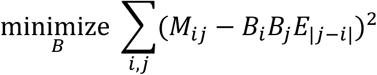

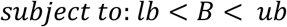

After obtaining the bias vector *B*, the normalized contact frequencies 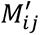 are calculated as 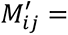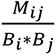

The bias vector *B* is initialized as the square root of the normalized one-dimensional coverage of the contact matrix:

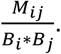

where *S*_𝑖_ = ∑*_j_* *M_ij_* represents the coverage of bin *i*, and 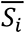 is the mean coverage across all loci.

Raichu employs the dual annealing optimization algorithm, with the L-BFGS-B algorithm as the local minimizer. To manage the computational demands of evaluating the objective function across millions or billions of data points, Raichu incorporates two performance-optimization strategies: (1) A sliding window [*s_i_, s_i_+w*] of size w is applied to each chromosome, with a 10% overlap between consecutive windows (*s_i+1_=s_i_*+0.9*w*). Bias vectors are computed independently for each window, and overlapping regions are averaged to produce a single bias vector for the chromosome. (2) The Numba library is used to enhance computational efficiency when evaluating the objective function.

Raichu provides several tunable parameters to enhance flexibility: (1) sliding window size (*w*), which is set to 200 bins by default; (2) maximum global search iterations for dual annealing (*m*), which is set to 100 iterations by default; and (3) bias vector search bounds (*lb, ub*), which are set to 0.001 and 1000 for Hi-C and single-cell Hi-C, and to 0.2 and 5 for HiChIP in this study.

### Hi-C, HiChIP, and ChIP-Seq data processing

All genomic datasets used in this study were mapped to the hg38 human reference genome. The processed Hi-C contact matrix for the GM12878 cell line (Figures 1-3), available at multiple resolutions in .mcool format, was downloaded from the 4DN data portal (accession code: 4DNFIXP4QG5B). Raw Hi-C reads for wild-type and engineered GM12878 cells (Figure 5) were obtained from the GEO database (accession code: GSE145997) and processed into contact matrices using the runHiC Python package (v0.9.0; https://pypi.org/project/runHiC/). Raw H3K27ac HiChIP reads for K562 cells (Figure 6) were downloaded from the GEO database (accession code: GSE101498) and processed into .mcool format using the same runHiC pipeline.

The H3K27ac and GATA3 ChIP-Seq peak files for wild-type and engineered GM12878 cells were also downloaded from GEO (accession code: GSE145997). Since the original peak files were mapped to hg19 genome coordinates, the coordinates were converted to hg38 using HiCLift^46^ (conversion script available at https://github.com/XiaoTaoWang/Raichu/tree/main/benchmark-analysis).

### Calculation of average contact signals between ChIP-Seq peaks of varying binding strength

For this analysis, the following procedures were followed:

1. TADs were identified using HiTAD^47^ (v 0.4.5-r1; https://xiaotaowang.github.io/TADLib/hitad.html) on GM12878 Hi-C contact maps at 25kb resolution. In the HiTAD output, only regions marked with “0” in the last column were classified as TADs, while all other regions were treated as sub-TADs.
2. ChIP-Seq peaks for CTCF, RAD21, POLR2A, and H3K27ac in GM12878 cells were downloaded from ENCODE (see Supplementary Table 2 for accession codes). Peaks for each factor were sorted and classified into eight groups based on binding strength, using the value in the 7th column of the downloaded peak file.
3. Average contact signals were calculated at 5kb resolution between each pair of peak groups. For bins containing multiple peaks, only the peak with the highest binding strength was considered. Only intra-TAD contacts were included, and contact signals were normalized by dividing the observed signal by the expected signal at the corresponding genomic distance (observed/expected).

### Loop detection in Hi-C data using HiCCUPS and Mustache

To benchmark the performance of Raichu in loop detection, we tested two tools: 1) A Python implementation of the original HiCCUPS algorithm^3^ (v0.3.8; https://github.com/XiaoTaoWang/HiCPeaks); and 2) Mustache^24^ (https://github.com/ay-lab/mustache). Our objective was to demonstrate that Raichu consistently enhances chromatin loop detection compared to existing normalization methods, regardless of the loop detection software used.

For HiCCUPS, the following parameters were applied: “--pw 1 2 4 --ww 3 5 7 --only-anchors -- maxapart 4000000”. For Mustache, the parameters were set as: “-pt 0.05”. Both tools were run on contact matrices at 5kb and 10kb resolutions separately. To generate a non-redundant list of detected loops for each tool, results from both resolutions were combined using the “combine-resolutions” command from the HiCPeaks Python package (v0.3.8; https://github.com/XiaoTaoWang/HiCPeaks) with the following parameters: “-G 10000 -M 100000 --max-res 10000”.

### Calculation of the overlap between two loop sets

For each loop coordinate (*i, j*) in loop set A, if there exists a loop (*i’, j’*) in loop set B such that the Euclidean distance between (*i, j*) and (*i’, j’*) is less than min (0.2×|*i-j*|, 50kb), we define (*i, j*) as preserved in *B*. Using this definition, we evaluated whether loops detected by Hi-C could be supported by orthogonal ChIA-PET/HiChIP interactions (Supplementary Table 2).

When generating the Venn diagram of loop sets obtained from different normalization methods, we applied an additional matching criterion: since a loop in one set might match multiple loops in another set based on the distance threshold, we ensured each loop matched only once by searching for the closest loop coordinate in the other set. This approach guaranteed a unique match for each loop meeting the distance criterion. The custom matching script is available at https://github.com/XiaoTaoWang/Raichu/tree/main/benchmark-analysis.

### Definition of promoter and enhancer regions

Promoter and enhancer regions in GM12878 cells were defined based on ChromHMM chromatin state annotations downloaded from the UCSC Genome Browser (https://genome.ucsc.edu/cgi-bin/hgTables). The annotation file was based on a 15-state ChromHMM model, which includes the following chromatin states: “1_Active_Promoter", “2_Weak_Promoter”, “3_Poised_Promoter”, “4_Strong_Enhancer”, “5_Strong_Enhancer”, “6_Weak_Enhancer”, “7_Weak_Enhancer”, “8_Insulator”, “9_Txn_Transition”, “10_Txn_Elongation”, “11_Weak_Txn”, “12_Repressed”, “13_Heterochrom/lo”, “14_Repetitive/CNV”, and “15_Repetitive/CNV”. Regions annotated as “1_Active_Promoter", “2_Weak_Promoter”, or “3_Poised_Promoter” were classified as promoters, while regions annotated as “4_Strong_Enhancer”, “5_Strong_Enhancer”, “6_Weak_Enhancer”, or “7_Weak_Enhancer” were classified as enhancers.

Since the original annotations were in hg19 coordinates, they were converted to hg38 using HiCLift^46^ (conversion script available at https://github.com/XiaoTaoWang/Raichu/tree/main/benchmark-analysis).

### Enrichment analysis of TFs and histone modifications

To explore the association of transcription factors (TFs) and histone modifications with each loop category (Fig. 2e, Supplementary Fig. 5d, and Supplementary Fig. 6d), we downloaded ENCODE ChIP-Seq peak files for 132 TFs and 10 histone modifications in GM12878 cells. For each TF or histone modification, we calculated both the fraction of loop anchors overlapping a ChIP-Seq peak and a fold-enrichment score.

Briefly, a non-redundant list of loop anchors was identified for each loop category. For each TF or histone modification, we iterated through this list and counted the number of anchors overlapping at least one ChIP-Seq peak. To calculate the fold enrichment score, we generated 100 random control sets by shuffling the loops and repeated the overlap calculation for each control set. Each random loop set preserved the genomic distance distribution between loop anchors and the number of loops per chromosome, ensuring that loop intervals did not overlap any gaps in the hg38 reference genome. The fold-enrichment score was then calculated by dividing the observed number of anchors overlapping ChIP-Seq peaks by the average number of overlaps in the random control sets.

For selected TFs and histone modifications, we also generated ChIP-Seq binding profiles around loop anchors. Specifically, the region spanning [-50kb, +50kb] around each loop anchor was divided into 50 bins of 2kb each. The number of ChIP-Seq peaks in each bin was counted, resulting in a 50- element array for each loop anchor. Each array was then normalized by dividing all values by the array’s mean value. Finally, the binding profile for each loop category was calculated by averaging the normalized values across all loop anchors at each bin position.

### Down-sampling of Hi-C data

To perform down-sampling at a rate of 𝛼 (0 < 𝛼 < 1), we used a binomial probability-based approach that does not require re-mapping of the raw reads. Specifically, for each non-zero pixel in the full contact matrix at 5kb resolution, the contact frequency was assigned as a random integer drawn from a binomial distribution with parameters *M_ij_* and *α*, where *M_ij_* represents the contact count in the full Hi-C matrix between bin 𝑖 and bin 𝑗. As both 5kb and 10kb contact matrices are required for loop detection (as described above), the “cooler coarsen” command was applied to the down-sampled 5kb contact matrices to generate the corresponding 10kb contact matrices.

### Single-cell Hi-C data analysis

The GM12878 single-cell Hi-C dataset in .pairs format was downloaded from GEO (accession code: GSE240128). Contact pairs were processed using HiCLift (https://github.com/XiaoTaoWang/HiCLift) to generate an .mcool file for each single cell, with the parameters “--output-format cool --in-assembly hg38 --out-assembly hg38”. Pseudo-bulk Hi-C maps were generated using the “cooler merge” command.

Loops were identified using pyHICCUPS (v0.3.8; https://github.com/XiaoTaoWang/HiCPeaks) at 5kb, 10kb, and 25kb resolutions individually, with the parameters ““--pw 1 2 4 --ww 3 5 7 --only-anchors -- maxapart 4000000”. Detected loops were then combined across resolutions using the “combine- resolutions” command from the HiCPeaks Python package (v0.3.8; https://github.com/XiaoTaoWang/HiCPeaks), with the parameters “-G 10000 -M 100000 --max-res 25000”.

To evaluate the precision of the detected loops, the union set of interactions compiled from orthogonal 3D genomic datasets (including Hi-C, CTCF ChIA-PET, RNAPII ChIA-PET, CTCF HiChIP, H3K27ac HiChIP, SMC1A HiChIP, H3K4me3 PLAC-Seq, TrAC-Loop, and HiCAR) was used as the reference (Supplementary Table 2). For recall evaluation, a high-confidence interaction set was defined as the reference, where each interaction was detectable in at least 3 of the 9 orthogonal datasets. F1 scores were calculated using the following equation:

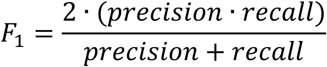

### Detection of differential loops

To detect differential loops between wild-type (C/C) and engineered (A/A) GM12878 cells, we applied the following approach to minimize instances where a loop is enriched in a contact map but not detected due to stringent p-value or fold-enrichment cutoffs (script available on GitHub at https://github.com/XiaoTaoWang/Raichu/tree/main/benchmark-analysis).

1. For each of the four contact maps (ICE-normalized and Raichu-normalized for wild-type and engineered GM12878 cells), we ran HiCCUPS using two parameter settings. For both settings, loops detected at 5kb and 10kb resolutions were merged using the “combine-resolutions” command, as described in the “Loop detection in Hi-C data using HiCCUPS and Mustache” section. The first setting used the default p-value and fold-enrichment cutoffs with parameters “-- pw 1 2 4 --ww 3 5 7 --only-anchors --maxapart 4000000”, which resulted in 8,803, 5,467, 11,114, and 6,562 loops for ICE-normalized maps in wild-type cells (*D_ICE, CC_*), ICE-normalized maps in engineered cells (*D_ICE, AA_*), Raichu-normalized maps in wild-type cells (*D_Raichu, CC_*), and Raichu- normalized maps in engineered cells (D*_Raichu, AA_*), respectively. The second setting used a more relaxed significance threshold with parameters “--pw 1 2 4 --ww 3 5 7 --only-anchors --maxapart 4000000 --siglevel 0.1 --sumq 0.05 --double-fold 1.5 --single-fold 1.75”, which resulted in 13,433, 8,688, 17,481, and 10,697 loops for ICE-normalized maps in wild-type cells (*L_ICE, CC_*), ICE-normalized maps in engineered cells (*L_ICE, AA_*), Raichu-normalized maps in wild-type cells (*L_Raichu, CC_*), and Raichu-normalized maps in engineered cells (*L_Raichu, AA_*), respectively.
2. The loop sets *D_ICE, CC_*, *D_ICE, AA_*, *D_Raichu, CC_*, and *D_Raichu, AA_* were defined as the final loops detected for the corresponding contact maps, while the loop sets *L_ICE, CC_*, *L_ICE, AA_*, *L_Raichu, CC_*, and *L_Raichu, AA_* were used to determine whether a loop from one set was potentially detectable in another contact map. For example, to check whether a loop (*i, j*) from *D_ICE, CC_* was detectable in the Raichu- normalized contact maps for wild-type GM12878 cells, we searched for a loop (*i’, j’*) in *L_Raichu, CC_* such that |*i-i’*| < min(0.2×|*i-j*|, 50kb) and |*j-j’*| < min(0.2×|*i-j*|, 50kb).

### Aggregate Peak Analysis

To evaluate the overall enrichment of chromatin loop signals in a contact map, we performed Aggregate Peak Analysis (APA). For a given list of chromatin loops or interactions, contact frequencies within w × w submatrices were extracted around the two-dimensional coordinates of each loop. The signals within each submatrix were normalized by dividing by their mean values. Next, submatrices with average signals above the 99th percentile or below the 1st percentile were filtered out to minimize biases from outliers. The average signal at each location was then calculated across all loops and visualized as heatmaps. In Figures 2h and 6g, contact matrices at 5kb resolution were used, with a window size (w) set to 15. In Figure 5d, contact matrices at 10kb resolution were used, with a window size of 11.

In Figures 2h and 5d, the z-score displayed on each APA plot was calcualted by comparing the signal at the center pixel with the signals in the lower-left 3 × 3 corner of the plot. In Figure 6g, the z-score was calculated by comparing the signal at the center pixel with the signals within the central 7 × 7 window.

### Aggregation analysis of TADs

To evaluate the enrichment of intra-TAD signals in the original, ICE-normalized, and Raichu-normalized H3K27ac HiChIP contact maps in K562 cells, we performed an aggregation analysis on regions surrounding TADs. To account for the distance-dependent decay of interaction frequencies, the interaction signal for each pixel was normalized by dividing it by the average signal across all bins at the same genomic distance.

For each TAD defined by the coordinates [𝑠_𝑖_, 𝑒_𝑖_], distance-normalized signals were calculated within the extended region [(3𝑠_𝑖_ − 𝑒_𝑖_)/2, (3𝑒_𝑖_ − 𝑠_𝑖_)/2], resulting in a matrix 𝑀_𝑖_. To aggregate TADs of varying sizes, the “resize” function from the scikit-image package (v0.20.0) was applied to resize each 𝑀_𝑖_ to a uniform shape of 60 × 60. Finally, all resized matrices were averaged to compute the mean signal at each location.

### Detection of significant interactions in HiChIP data

As a proof of concept, we focused on detecting significant interactions on chromosome 21 in K562 cells, as this chromosome is relatively free of structural variations (SVs) in this cancer cell line. The absence of SVs is critical, because their presence can distort interaction frequencies and compromise the accuracy of significant interaction detection^48, 49^.

We applied an algorithm similar to HiCCUPS to identify statistically significant interactions from ICE- normalized and Raichu-normalized H3K27ac HiChIP contact maps at 5kb resolution. Significant interactions were determined by comparing observed interaction frequencies against both global and local interaction backgrounds.

Specifically, the global interaction background 𝐸_𝑘_ for interactions at a genomic distance 𝑘 was calculated as 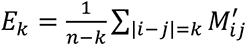, where 𝑛 is the total number of bins on chromosome 21 at 5kb resolution, and 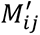 represents the normalized interaction frequency. Two local background filters, the donut filter 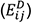 and the lower-left filter 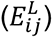, were applied following the definitions in the original HiCCUPS algorithm.

### Donut filter

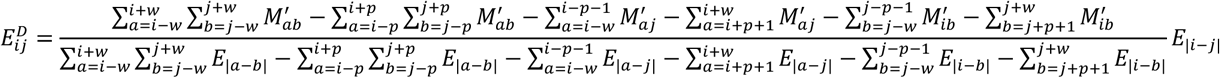

Lower-left filter:

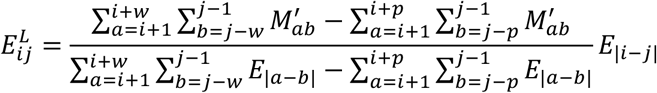

Parameters 𝑝 and 𝑤, which define the interaction peak and background window sizes, were set to 4 and 7, respectively.

Interactions with at least 25 total read counts within a 15 × 15 bin window and spanning genomic distances no greater than 3Mb were considered. For each interaction (𝑖, 𝑗), statistical significance was evaluated using a Poisson distribution with 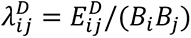 or 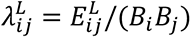, where 𝐵_𝑖_ and 𝐵_*j*_ are bias factors. Two p-values 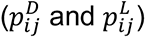 were computed and converted to q-values 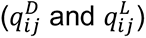 using the “multipletests” function (method=’fdr_bh’) from the statsmodels Python package.

During the q-value calculation, interactions were grouped based on their biases using a “λ-chunking” procedure. Specifically, for each interaction (𝑖, 𝑗), a bias value 𝐵_𝑖j_ = 𝐵_𝑖_𝐵_*j*_ was assigned. Interactions with *B_ij_* < 1 were grouped together, while subsequent groups were spaced logarithmically (every 2^1/3^). The “multipletests” function was then applied independently to each group. For downstream analysis, interactions on chromosome 21 were ranked by q-values, and the top 4,000, 3,000, 2,000, and 1,000 interactions were selected.

To detect significant interactions directly from raw contact maps without normalization, the global background 𝐸_𝑘_ was calculated as 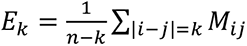, and local backgrounds (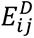 and 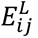) were computed using the raw contact matrix (𝑀) instead of the normalized matrix (𝑀^′^). Additional adjustments included: (1) assigning 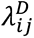 and 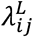 as 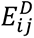 and 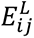, respectively; and (2) calculating q-values without bias grouping (all interactions treated as a single group). All other steps followed the workflow described for normalized data.

### Identification of high-confidence H3K27ac enriched regions

To evaluate the precision and recall of the original, ICE-normalized, and Raichu-normalized H3K27ac HiChIP signals in detecting enhancer-associated interactions, we compiled a list of high-confidence H3K27ac-enriched regions using the following steps:

1. H3K27ac ChIP-Seq peaks were downloaded from the ENCODE data portal (accession code: ENCFF544LXB) and designated as set A.
2. StripeCaller (v0.1.0; https://github.com/XiaoTaoWang/StripeCaller) was applied to H3K27ac HiChIP contact maps at 5kb and 10kb resolutions using the parameter “--use-raw”. Stripe anchors detected at 5kb and 10kb resolutions were designated as sets ***B*** and ***C***, respectively.
3. Each region in sets ***B*** and ***C*** was iterated and checked for overlap with regions in the other sets. A region was included in the final list of high-confidence H3K27ac-enriched regions if it appeared in at least two of the sets (***A, B***, or ***C***).

### Enrichment analysis of chromatin states

To generate Figure 6e, ChromHMM chromatin state annotations for K562 cells were downloaded from the UCSC Genome Browser (https://genome.ucsc.edu/cgi-bin/hgTables) and converted from hg19 to hg38 coordinates using HiCLift (conversion script available on GitHub at https://github.com/XiaoTaoWang/Raichu/tree/main/benchmark-analysis).

To characterize the chromatin states of the anchors in a given list of HiChIP interactions, we calculated a fold-enrichment score for each state by comparing the overlap of anchors with real ChromHMM annotations to overlaps with 100 randomly shuffled control annotations.

Specifically, for each chromatin state, we iterated through the interaction anchor list and counted the number of anchors overlapping at least one region annotated with that state. We then randomly shuffled the regions for that state across the genome to create 100 control sets and repeated the overlap calculation for each control. For each control, we maintained the size distribution and number of random regions per chromosome to match the real ChromHMM annotations, ensuring that no region intervals overlapped gaps in the hg38 reference genome. Finally, the fold-enrichment score was calculated by dividing the number of anchors overlapping a specific chromatin state by the average number of overlaps in the random control sets for that state.

## Data availability

Details of all datasets analyzed in this study are summarized in Supplementary Table 2.

## Code availability

The Raichu source code is publicly available on GitHub at https://github.com/XiaoTaoWang/Raichu/, and the scripts used for data analysis can be accessed at https://github.com/XiaoTaoWang/Raichu/tree/main/benchmark-analysis.

## Supporting information

Supplementary information including 11 supplementary figures and 1 supplementary table

## Acknowledgements

This work was supported by the National Natural Science Foundation of China (No. 32470698), the National Key Research and Development Program of China (No. 2022YFC2703400), the CAMS Innovation Fund for Medical Sciences (No. 2019-I2M-5-064), and Shanghai Frontiers Science Research Center of Reproduction and Development.

## Author contributions

X.W. conceived the project, implemented the Raichu framework, and led the data analysis. D.S. co- led the data analysis. F.X. and H.Y. contributed to the data analysis, with F.X. also assisting in figure design. Y.L. supported data collection. And L.J. assisted with software testing.

