## Supplementary information including 11 supplementary figures and 1 supplementary table for "Boosting the detection of enhancer-promoter loops via novel normalization methods for chromatin interaction data"

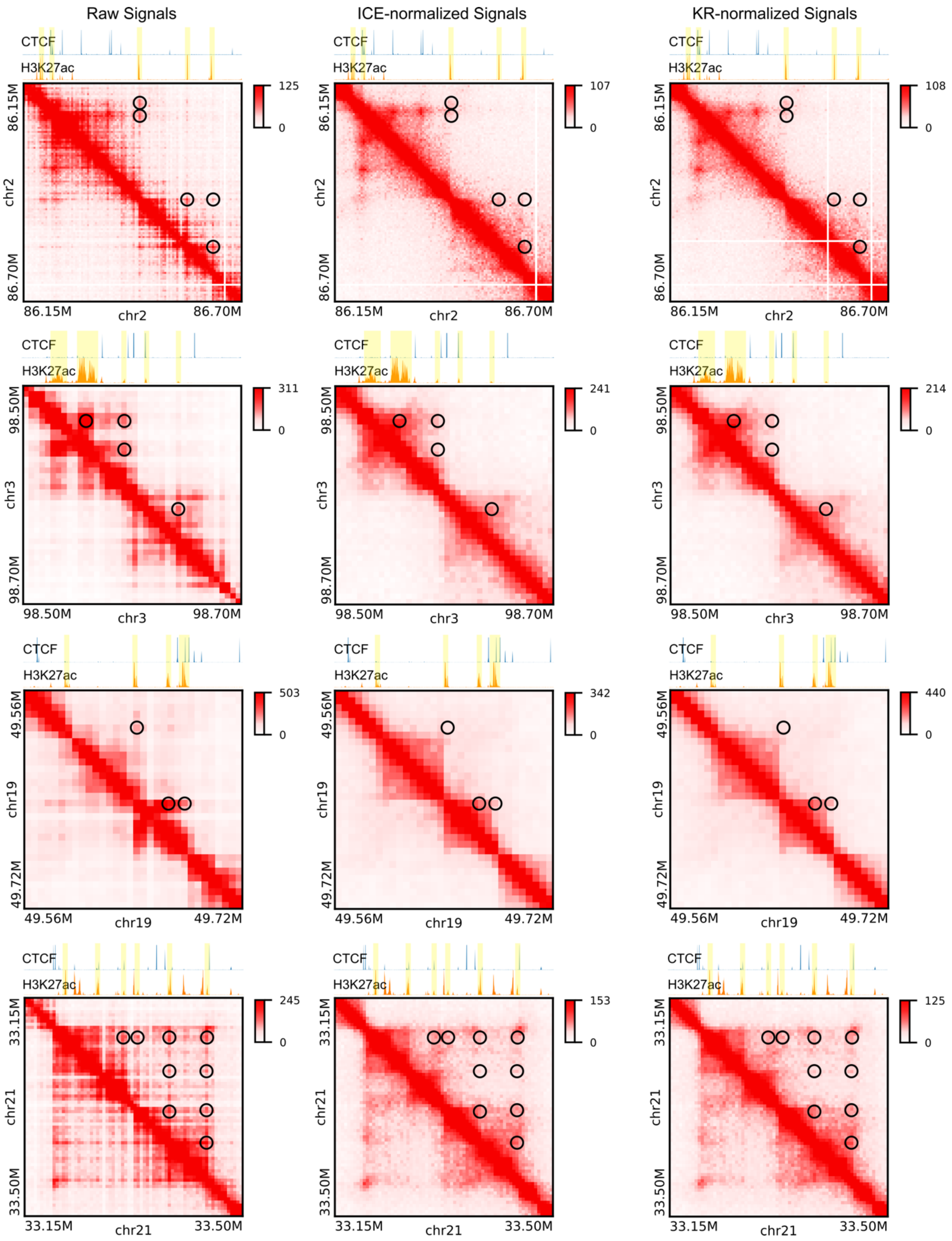


**Supplementary Figure 1. Examples illustrating the depletion of transcription-related loops after ICE or KR normalization.** In each row, raw signals, ICE-normalized signals, and KR-normalized signals are compared for the same region in GM12878 cells. Black circles indicate loops that are strongly enriched in the raw Hi-C map but become undetectable after ICE or KR normalization. Above the Hi-C heatmaps, CTCF and H3K27ac ChIP-seq tracks are displayed, with yellow bars marking the positions of the loop anchors corresponding to the loops highlighted by the black circles.

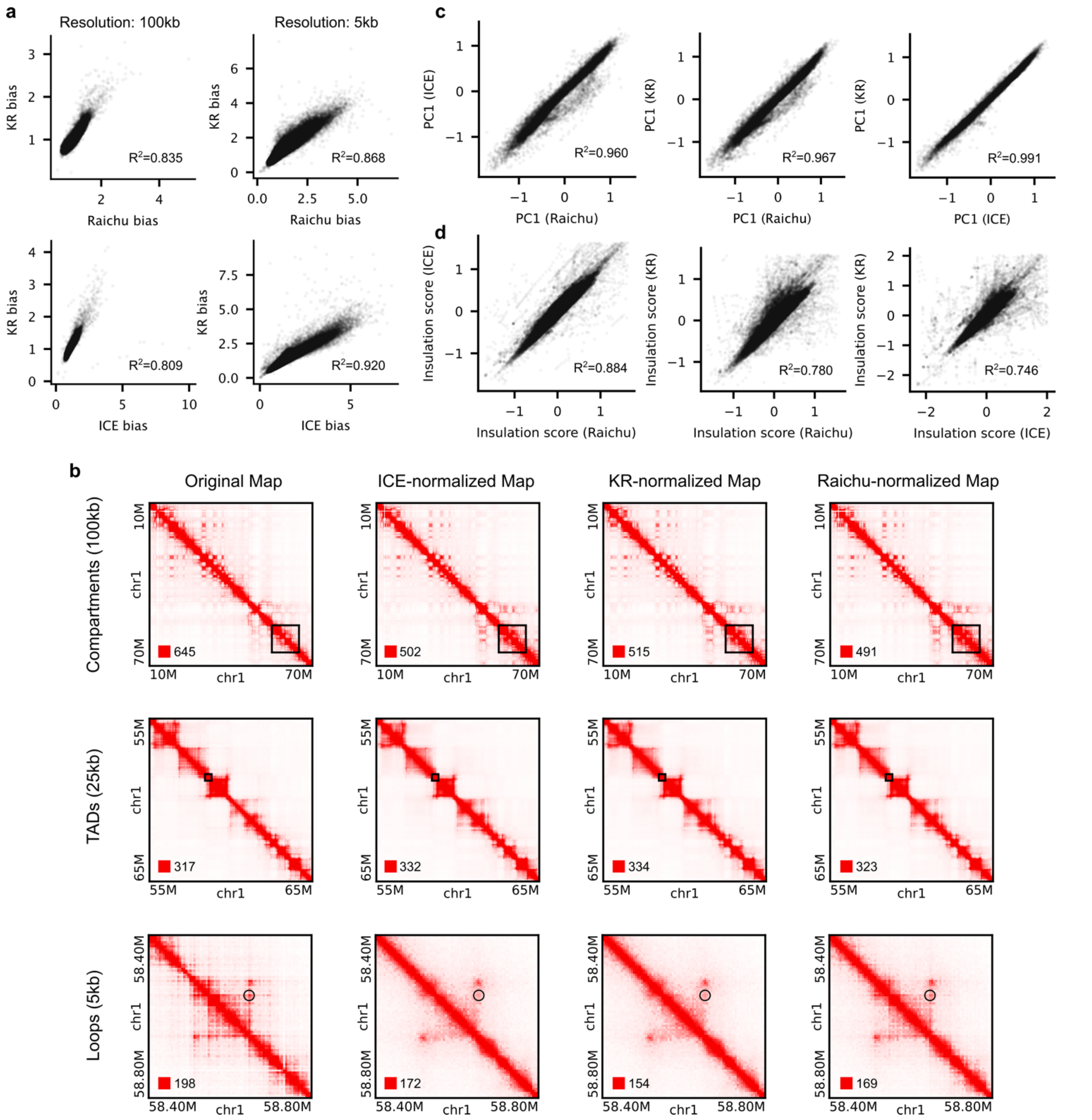


**Supplementary Figure 2. Raichu is comparable to existing Hi-C data normalization methods in detecting higher-order chromatin organization features. a,** (First row) Comparison of bias vectors calculated by Raichu and KR at 100kb and 5kb resolutions. (Second row) Comparison of bias vectors calculated by ICE and KR at 100kb and 5kb resolutions. **b**, Example regions showing raw contact signals, ICE-normalized signals, KR-normalized signals, and Raichu-normalized signals at different resolutions. **c**, Comparison of the first principal component (PC1), which characterizes chromatin compartment patterns, at 100kb resolution across different normalization methods. **d**, Comparison of insulation scores, which characterize topologically associating domain (TAD) patterns, at 25kb resolution across different normalization methods.

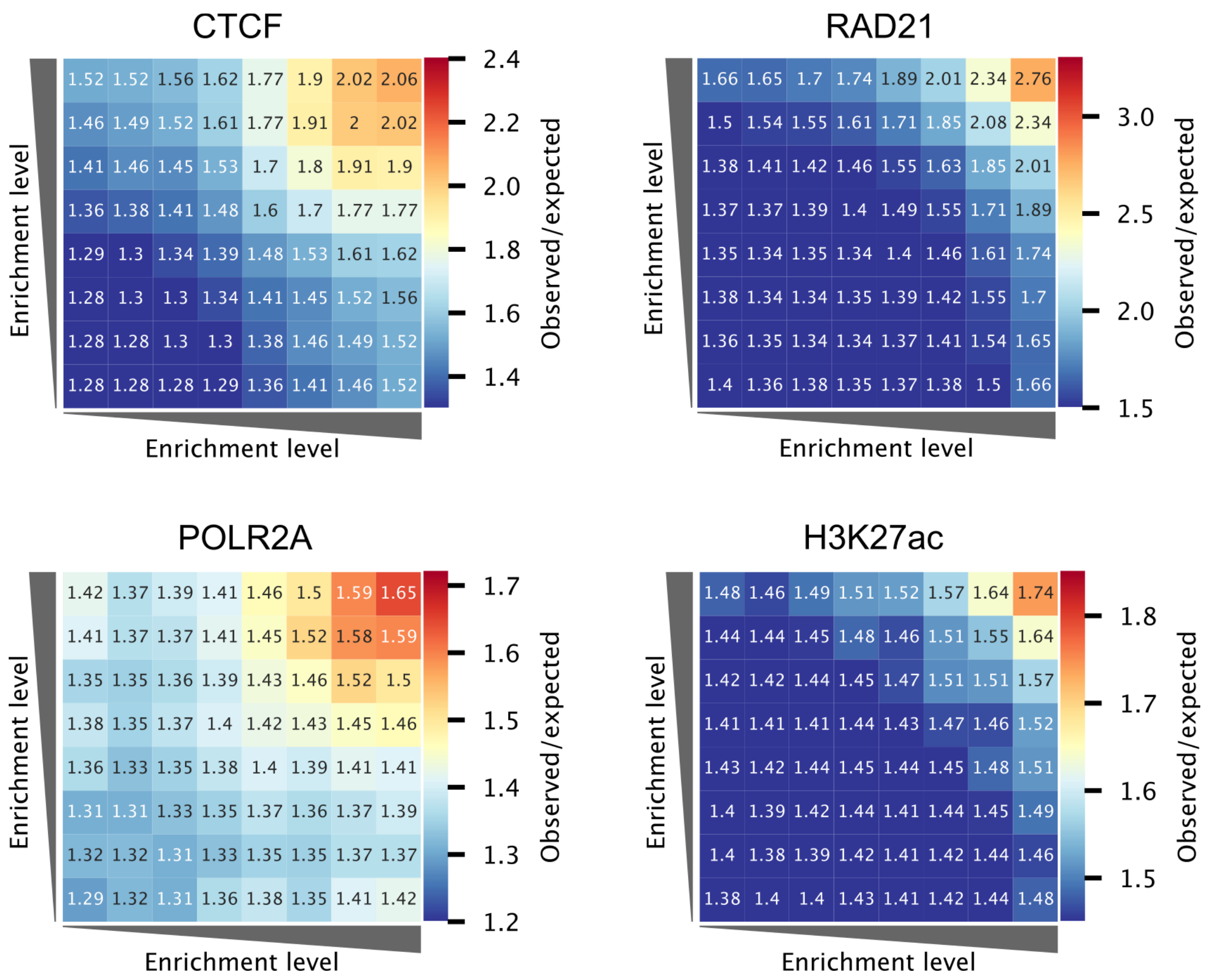


**Supplementary Figure 3. Average (observed/expected) KR-normalized contact signals between binding peaks of the indicated transcription factors or histone modifications.** Similar plots for ICE- and Raichu-normalized signals are presented in Figure 1 of the main text.

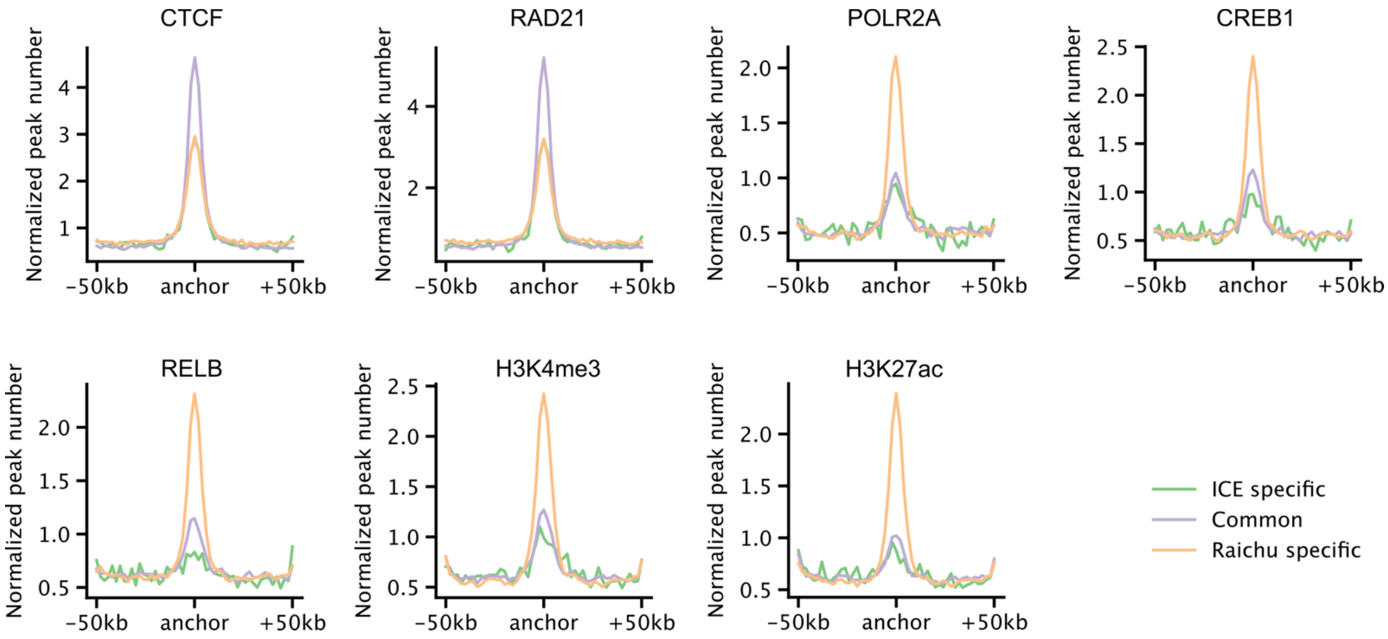


**Supplementary Figure 4. ChIP-Seq binding profiles of selected transcription factors and histone modifications around loop anchors.** Different colors indicate different loop categories.

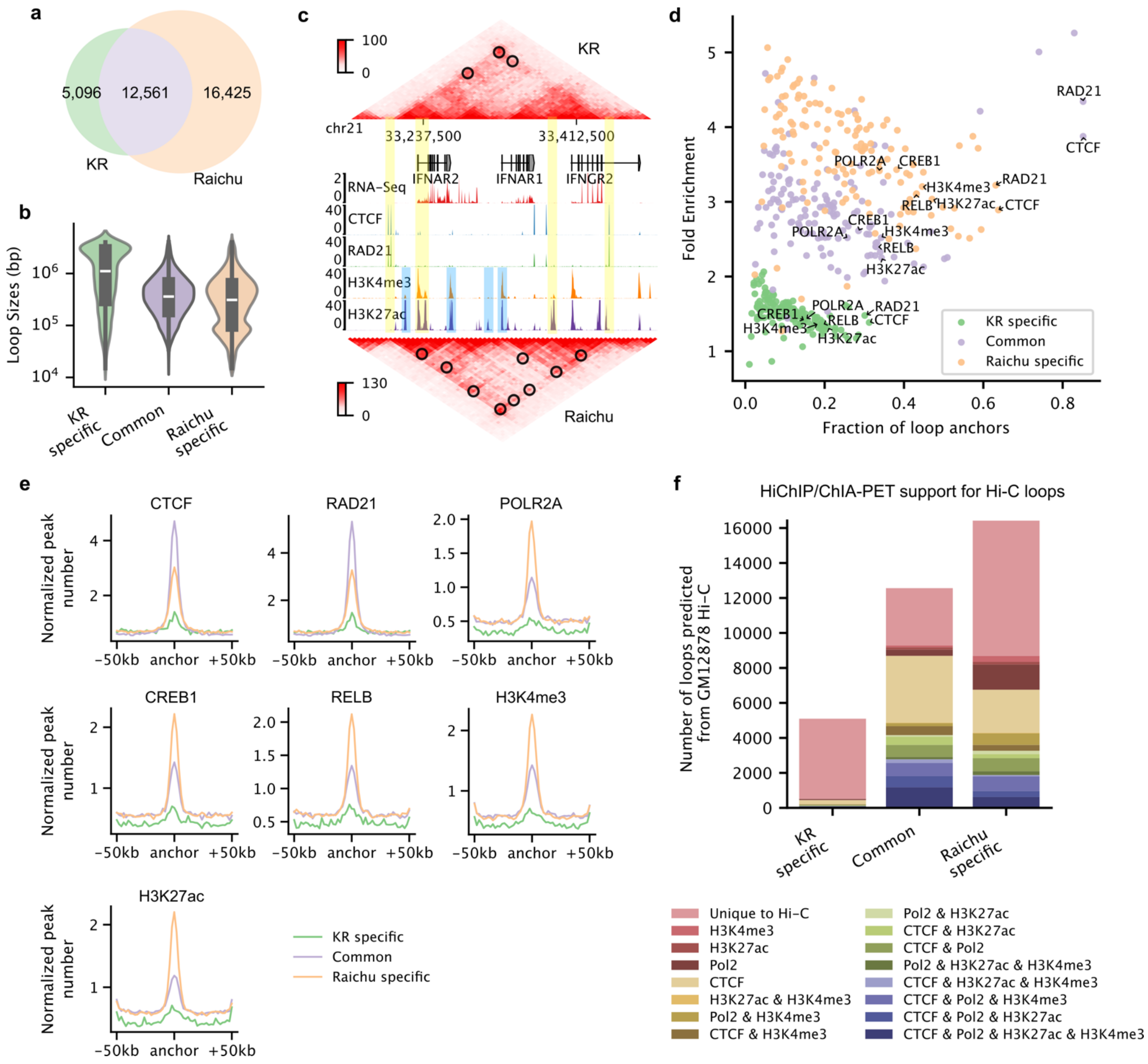


**Supplementary Figure 5. Raichu identifies thousands of transcription-related loops missed by KR.** HiCCUPS was used for loop detection. **a,** Venn diagram showing the overlap of loops detected by KR and Raichu in GM12878 cells. **b**, Violin plots comparing the sizes of KR-specific, Raichu-specific, and common loops detected by both KR and Raichu. **c**, Comparison of contact signals for an example region between KR and Raichu. Contact heatmaps are shown alongside gene annotations, RNA-seq data, and ChIP-seq signals for selected transcription factors and histone modifications. Black circles mark identified loops, yellow bars highlight common loop anchors detected by both KR and Raichu, and blue bars highlight Raichu-specific loop anchors. **d,** Fraction of loop anchors bound versus fold enrichment for 132 transcription factors and 10 histone modifications, with colors representing the loop categories from the Venn diagram in panel a. **e**, ChIP-seq binding profiles of selected transcription factors and histone modifications around loop anchors for different loop categories. **f**, Overlap with orthogonal ChIA-PET/HiChIP interactions for different loop categories.

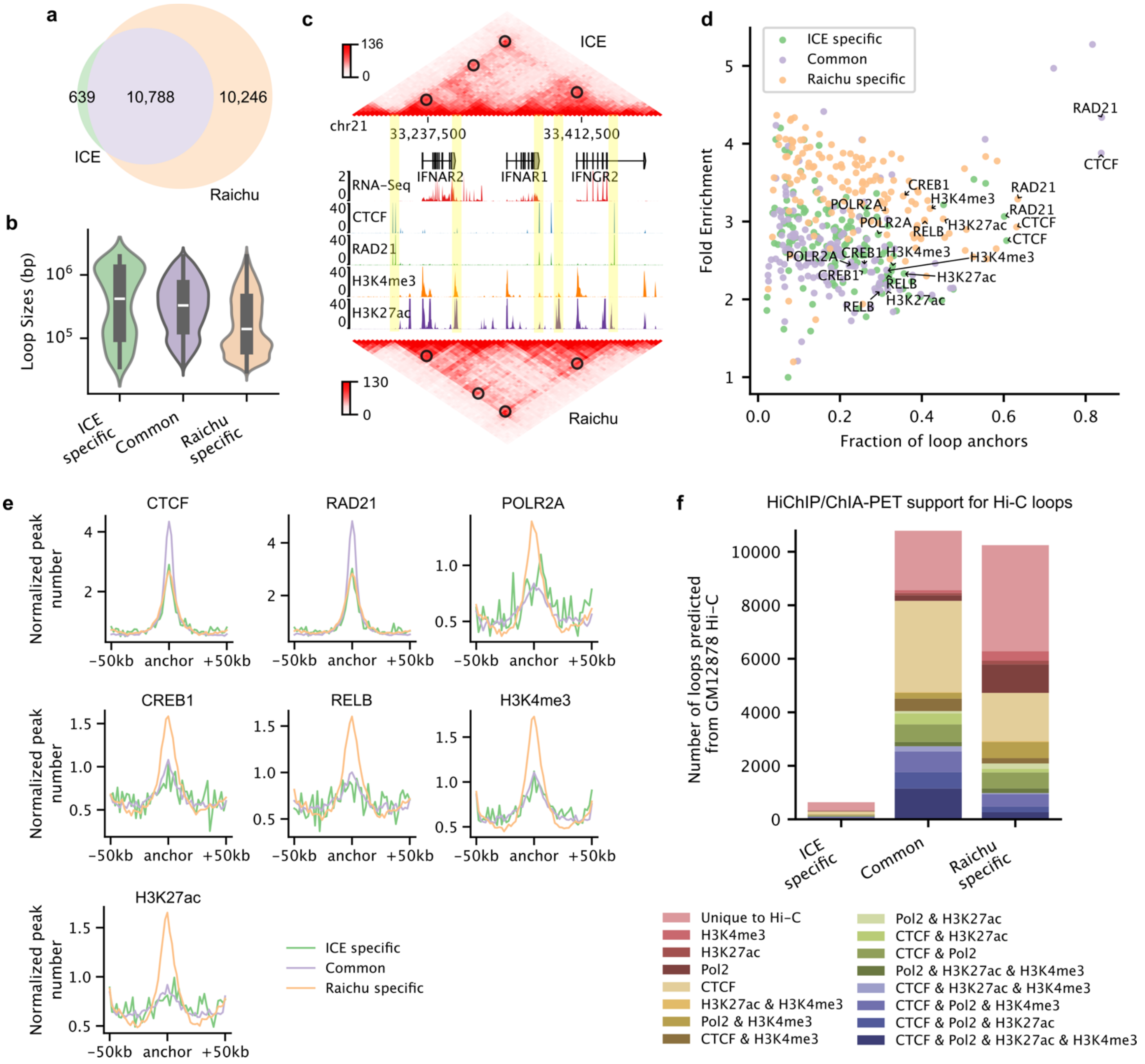


**Supplementary Figure 6. Raichu identifies thousands of transcription-related loops missed by ICE.** Mustache was used for loop detection. **a,** Venn diagram showing the overlap of loops detected by ICE and Raichu in GM12878 cells. **b**, Violin plots comparing the sizes of ICE-specific, Raichu-specific, and common loops detected by both ICE and Raichu. **c**, Comparison of contact signals for an example region between ICE and Raichu. Contact heatmaps are shown alongside gene annotations, RNA-seq data, and ChIP-seq signals for selected transcription factors and histone modifications. Black circles denote identified loops on the corresponding maps. **d,** Fraction of loop anchors bound versus fold enrichment for 132 transcription factors and 10 histone modifications, with colors representing the loop categories from the Venn diagram in panel a. **e**, ChIP-seq binding profiles of selected transcription factors and histone modifications around loop anchors for different loop categories. **f**, Overlap with orthogonal ChIA-PET/HiChIP interactions for different loop categories.

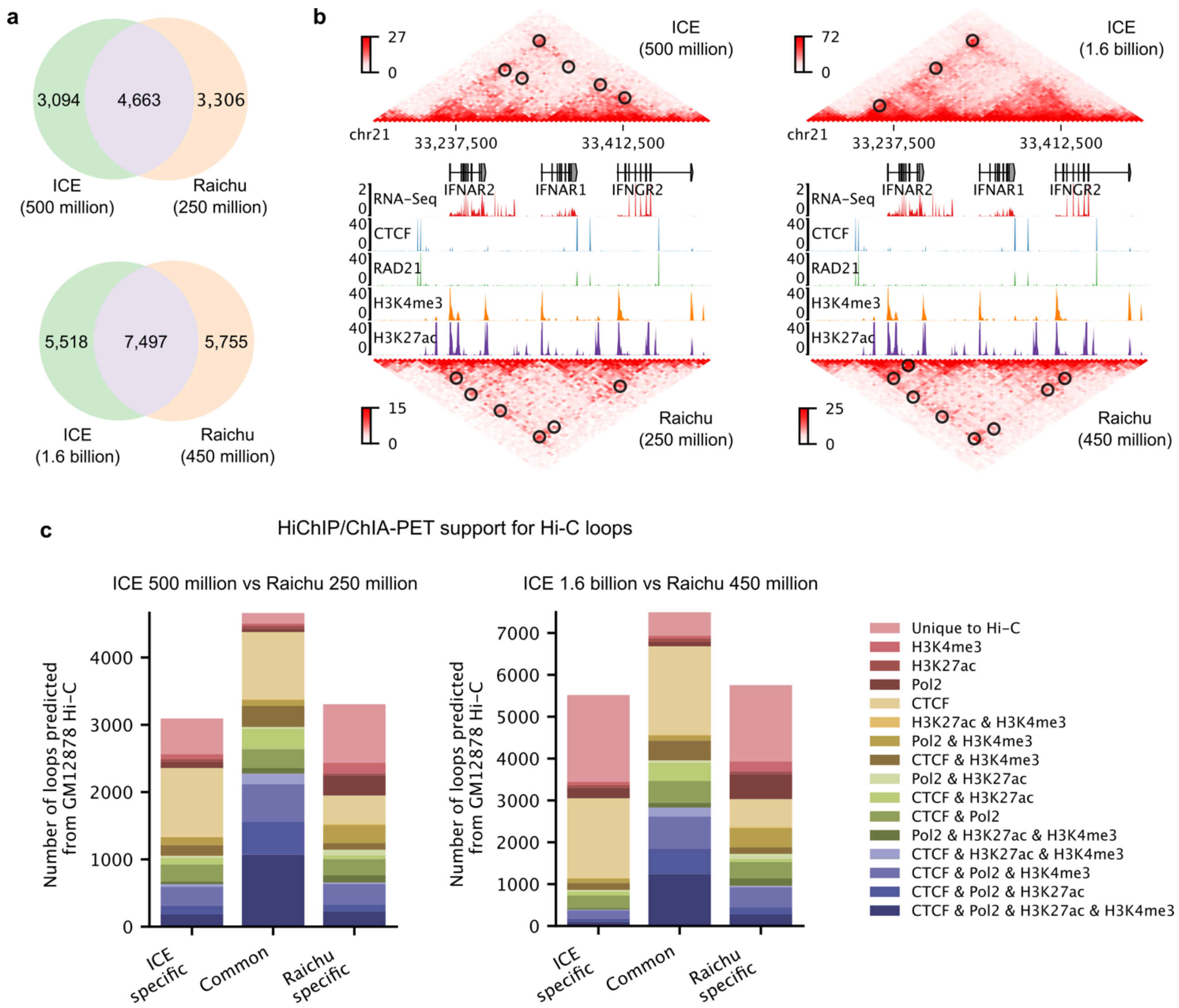


**Supplementary Figure 7. Raichu performs well across different sequencing depths.** HiCCUPS was used for loop detection. **a,** Venn diagrams comparing loops detected by ICE with 500 million usable reads to those detected by Raichu with 250 million usable reads, and loops detected by ICE with 1.6 billion usable reads to those detected by Raichu with 450 million usable reads. **b**, Comparison of loops identified by ICE and Raichu at the indicated sequencing depths. Contact heatmaps, gene annotations, RNA-Seq data, and ChIP-Seq signals for selected transcription factors and histone modifications are shown. Black circles denote identified loops. **c**, Overlap with orthogonal ChIA-PET/HiChIP interactions for ICE-specific, Raichu-specific, and common loops detected by both ICE and Raichu, as defined in panel a.

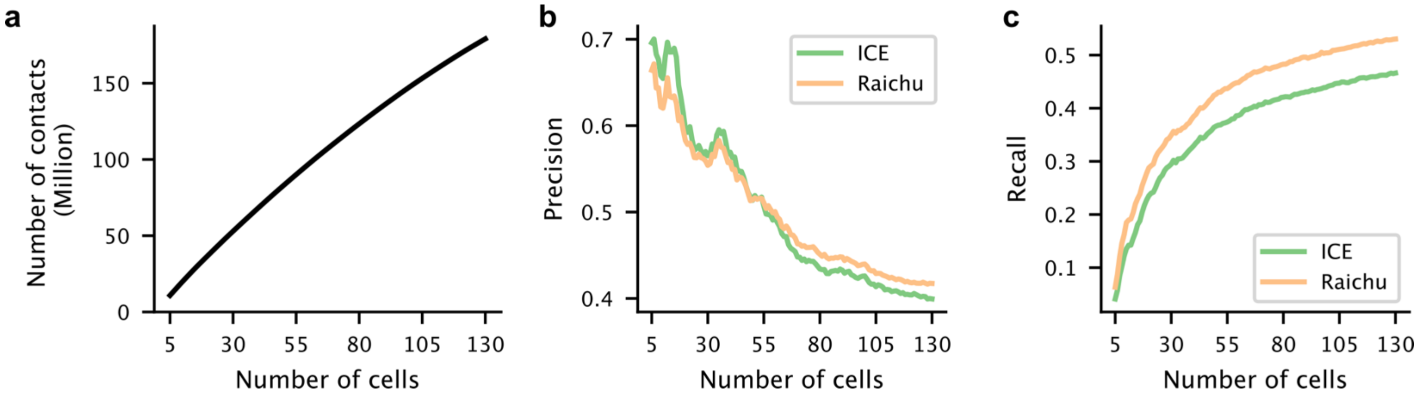


**Supplementary Figure 8. Raichu outperforms ICE in loop detection for single-cell Hi-C data. a,** Number of chromatin contacts as a function of the number of merged GM12878 single cells. **b-c**, Precision and recall of loops detected by ICE and Raichu across varying numbers of merged GM12878 cells.

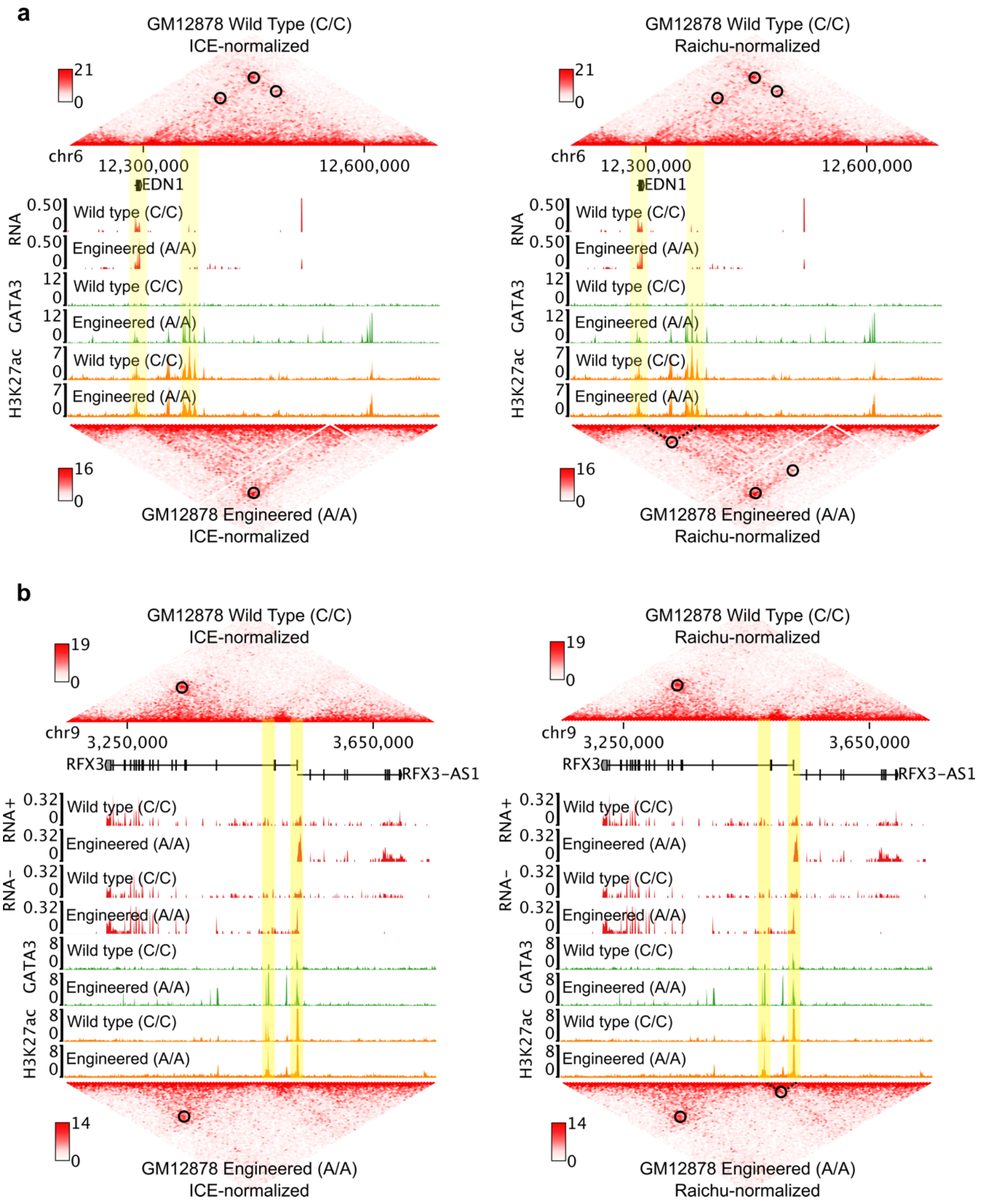


**Supplementary Figure 9. Examples illustrating Raichu's ability to detect unique A/A-specific loops associated with target gene activation.**

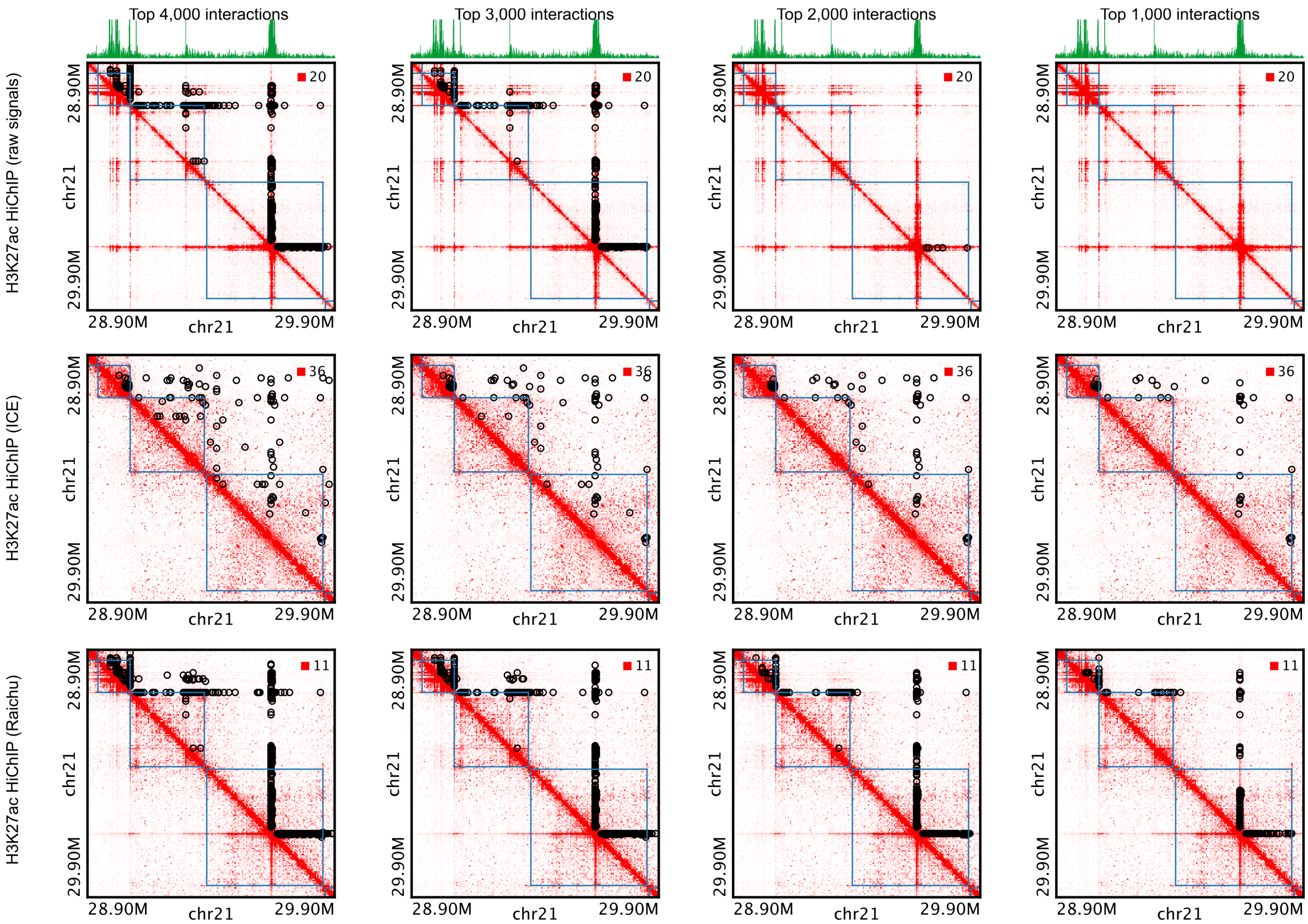


**Supplementary Figure 10. Example comparing different normalization methods for H3K27ac HiChIP data in K562 cells.** The top green track represents the one-dimensional coverage of the HiChIP data, with peaks indicating H3K27ac-enriched regions. Blue squares denote TAD regions identified from Hi-C data in the same cell line. Black circles mark the significant interactions identified. For each normalization method, interactions were ranked by p-values, and a specified number of top interactions are displayed to illustrate the sensitivity and specificity of each method.

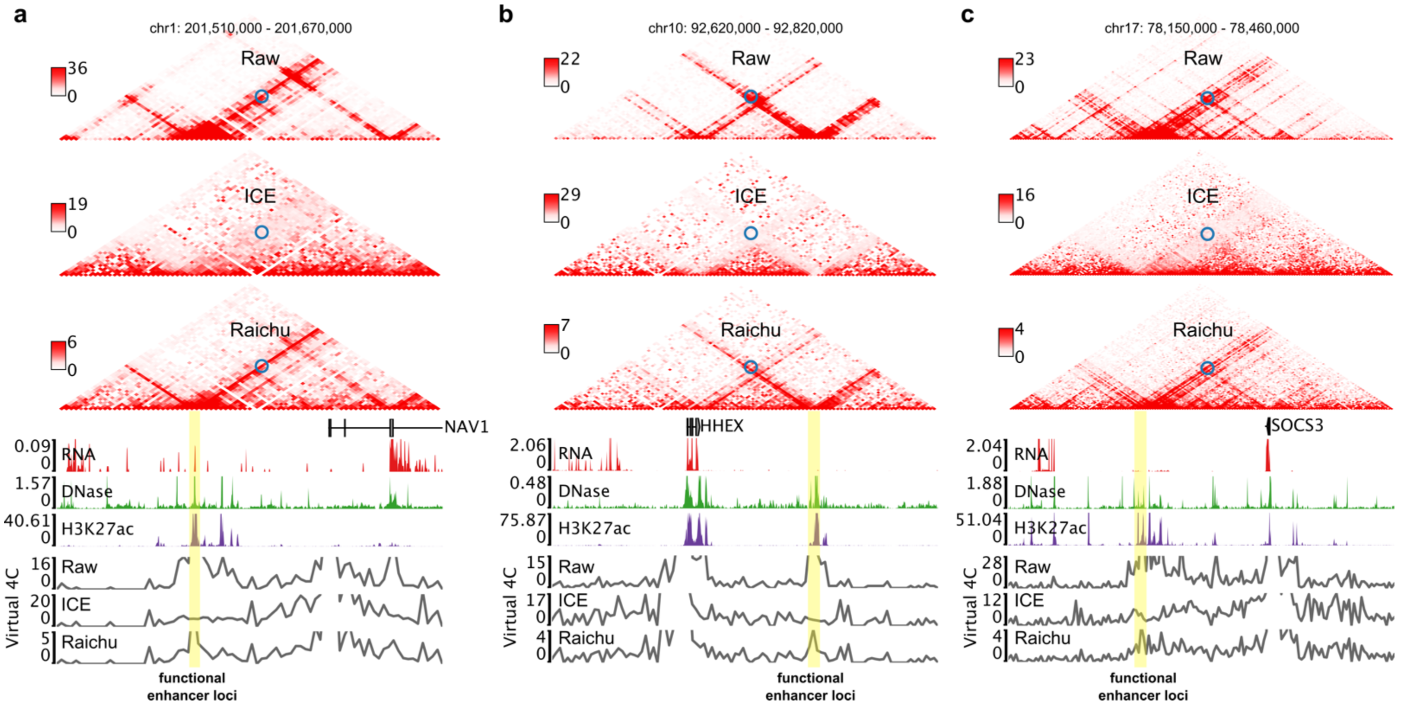


**Supplementary Figure 11. Examples comparing different normalization methods for H3K27ac HiChIP data in detecting known functional enhancer-promoter interactions in K562 cells.** In each example, the yellow bar marks the enhancer, and blue circles indicate the functional interaction.

**Supplementary Table 1. Running time and memory usage of Raichu**

| Resolution | Window Sizes | Running with 4 processes | | Running with 8 processes | |
|---|---|---|---|---|---|
| | | Memory Usage | Running Time | Memory Usage | Running Time |
| 100kb | 200 | 1.6G | 0:19:47 | 2.8G | 0:10:08 |
| | 400 | 1.6G | 0:52:54 | 2.9G | 0:29:15 |
| 50kb | 200 | 2.6G | 0:43:34 | 4.0G | 0:24:48 |
| | 400 | 2.6G | 2:02:15 | 4.2G | 1:06:14 |
| 10kb | 200 | 14.4G | 2:13:30 | 18.6G | 1:13:07 |
| | 400 | 12.7G | 7:31:55 | 18.5G | 3:28:17 |
| 5kb | 200 | 17.0G | 4:04:11 | 26.6G | 2:33:47 |
| | 400 | 14.1G | 11:46:55 | 25.7G | 6:11:18 |

Test dataset: GM12878 (Rao 2014, ~4.01 billion filtered reads)

CPU: Hygon C86 7285 32-core Processor

Running time format: hrs: min: sec

The memory usage was recorded using Memory Profiler (https://github.com/pythonprofilers/memory_profiler) at 3-second intervals during the execution of Raichu, and the peak usage is shown in the table above.
